# Multifaceted B-cell response to transient HIV viraemia in elite controllers

**DOI:** 10.1101/2024.12.28.630596

**Authors:** Luke Muir, Ondrej Suchanek, Peter Thomas, Sarah A. Griffith, Emma Touizer, Chloe Rees-Spear, Masahiro Yoshida, Christopher L. Pinder, Marko Z. Nikolic, Katie J. Doores, Marit J. Van Gils, Ravindra K. Gupta, Menna R. Clatworthy, Laura E. McCoy

**Affiliations:** Institute for Immunity and Transplantation, Division of Infection and Immunity, University College London, UK; Molecular Immunity Unit, Department of Medicine, Medical Research Council Laboratory of Molecular Biology, University of Cambridge, Cambridge, UK; Cambridge University Hospitals NHS Foundation Trust, and NIHR Cambridge Biomedical Research Centre, Cambridge, UK; UCL Respiratory, Division of Medicine, University College London, London, UK; Cellular Genetics, Wellcome Sanger Institute, Cambridge, UK; Department of Infectious Diseases, School of Immunology & Microbial Sciences, King’s College London, London, UK; Amsterdam UMC, University of Amsterdam, Department of Medical microbiology and infection prevention, Amsterdam, the Netherlands; Amsterdam Instittute for Infection and Immunity, Infectious Diseases, Amsterdam, the Netherlands; Cambridge Institute of Therapeutic Immunology and Infectious Disease (CITIID), Cambridge, UK; Department of Medicine, University of Cambridge, Cambridge, UK

## Abstract

Chronic HIV infection drives B-cell dysfunction associated with the accumulation of tissue-like memory (TLMs) and activated memory B cells (MBCs) but decline in resting memory B cells. TLMs express multiple inhibitory receptors and lack response to soluble antigens. However, their origin and the mechanisms driving their expansion in HIV infection remain unclear. By using bulk BCR sequencing of memory B cell subsets from elite HIV controllers and an ART-controlled cohort, we revealed that TLMs (CD21^-^ CD27^-^ B cells) were significantly less mutated but also less diverse than other MBCs, suggesting an enrichment for innate-like B cells or that they belong to a less mature subset. Subsequent detailed multi-omics study of an immune response in an elite controller to a transient HIV viraemia demonstrated a functional increase in Env-reactive IgG and MBCs with non-TLM phenotype. Single-cell RNA/BCR sequencing of PMBCs enriched for B cells revealed an orchestrated TNF-alpha response followed by interferon alpha and gamma response across all B cells subsets. We noted an emergence of innate-like extrafollicular PLD4+ plasmablasts and previously undepreciated heterogeneity of a stable TLM population. We identified two distinct TLM subsets: TLM1 (T-bet^low^) and TLM2 (T-bet^hi^) that differed in their differentiation stage, isotype use and mutational burden. Surprisingly, both subsets (TLM1 more than TLM2) were enriched for IGHV4-34 segment use (associated with self-reactivity) and displayed a persistent activation and BCR signalling signature, indicating (with other BCR indices) a strong presence of innate-like B cells. However, pseudotime analyses revealed that TLMs contained not only innate-like B cells but also cells from the conventional memory B cell lineages, further highlighting the complexity of the whole TLM compartment. This study provides a new insight into multifaceted functional B-cell response to transient HIV viraemia as likely happens during the early phase of anti-retroviral therapy cessation, highlighting the TLM heterogeneity and the contribution of innate-like B cells which might have important clinical implications for anti-HIV vaccine and therapy design.

## INTRODUCTION

B cell dysfunction in people living with human immunodeficiency virus (HIV) (PLWH) has been widely described (Moir and Fauci 2009) and is associated with suboptimal B cell responses to both infection and vaccination (Malaspina, Moir et al. 2005, Titanji, De Milito et al. 2006, Touizer, Alrubayyi et al. 2023). Although antiretroviral therapy (ART) has proven effective in reducing the viral load in PLWH to undetectable levels and generally restores CD4^+^ T cell counts, B cell homeostasis is never fully restored even in those who have maintained ART and supressed viral loads for multiple years (Morris, Binley et al. 1998, Moir, Buckner et al. 2010, Pensieroso, Galli et al. 2013). Some of the key changes associated with HIV-mediated B cell dysfunction include hypergammaglobulinemia and the dysregulation of certain B cell subsets, including increased numbers of immature transitional B cells, activated memory B cells, tissue-like memory B cells (TLMs) and plasmablasts, whilst resting memory B cell numbers are typically reduced (Moir and Fauci 2009).

TLMs were first described in 2008 by Moir et al. and can account for >50% of the memory B cell population in untreated chronic HIV infection compared to ∼5% in HIV-negative individuals (Moir, Ho et al. 2008). Their name refers to their enriched expression of surface FcRL4, similar to the tissue B cell population previously identified in human tonsils (Ehrhardt, Hsu et al. 2005). TLMs lack expression of the typical memory B cell markers CD21 and CD27 and are CD10^-^ CD19^Hi^ CD20^Hi^ (Moir, Ho et al. 2008, Muellenbeck, Ueberheide et al. 2013). They have also been shown to be enriched for expression of the Th1-associated transcription factor T-bet and the integrin CD11c (Moir, Ho et al. 2008, Knox, Buggert et al. 2017). However, the most defining feature of this subset is their high expression of a variety of inhibitory receptors (alongside FcRL4), including CD22, CD85j and CD72 (Moir, Ho et al. 2008, Knox, Buggert et al. 2017). Further, TLMs typically display a chemokine receptor profile suggestive of homing to sites of inflammation, with increased expression of CXCR3 and CCR6 (Moir, Ho et al. 2008, Knox, Buggert et al. 2017). Functional analysis of TLMs initially indicated that they were significantly less responsive to soluble stimuli and appeared to have undergone fewer rounds of proliferation *in vivo* compared to classical memory B cells, resulting in the hypothesis that these were an exhausted B cell subset driven by chronic antigen exposure during infection (Moir, Ho et al. 2008). This hypothesis was further supported by the evidence that a knockdown of inhibitory receptor expression (including FCRL4, SIGLEC6 and CD85J) on these subsets using siRNA could partially restore the ability of TLMs to respond to soluble stimuli (Kardava, Moir et al. 2011). However, recent data has suggested that TLMs can respond to complex antigens, suggesting they may not be as functionally inert as first thought (Ambegaonkar, Kwak et al. 2020).

Expanded populations analogous to TLMs have been described in various other chronic inflammatory settings, such as Hepatitis B infection, Malaria infection, and systemic lupus erythematosus (SLE) (Portugal, Tipton et al. 2015, Burton, Pallett et al. 2018, Jenks, Cashman et al. 2020). Although a different label is used for this B cell subset in each of these conditions (e.g., “atypical” B cells in malaria or “double negative” B cells in SLE), the cells appear to share several hallmark characteristics, including the expression of inhibitory receptors, T-bet and a CD21^lo^ memory B cell phenotype. Malaria-associated atypical memory B cells are amongst the most studied, they expand during chronic infection forming up to 51% of total mature circulating B cells. Extensive scRNA sequencing analysis on TLM, atypical and DN cells from HIV, Malaria and SLE respectively, revealed remarkably similar transcriptional profiles for these clusters (Holla, Dizon et al. 2021). Overall, this suggests that there is a potential common mechanism behind the induction and expansion of these cells during chronic inflammation arising from different disease states.

To date the mechanisms driving the development of TLM/atypical B cells have not yet been fully defined. Recent data from murine models has shown that the IFN-γ dynamics associated with chronic LCMV infection can induce an epigenetically distinct B cell subset (Cooper, Xu et al. 2024) whilst data from individuals who acquired malaria suggested an important role for IFN-γ (Obeng-Adjei, Portugal et al. 2017, Ambegaonkar, Nagata et al. 2019, Holla, Dizon et al. 2021). In contrast, a previous study in PLWH with uncontrolled viraemia, found the majority of HIV envelope glycoprotein (Env)-reactive memory B cells were T-bet^+^, a marker which has been linked to TLMs, suggesting that there may be a role for antigen-driven effects (Moir, Ho et al. 2008, Kardava, Moir et al. 2014, Knox, Buggert et al. 2017, Burton, Pallett et al. 2018). However, as TLMs can account for ∼50% of the class-switched memory B cell population during chronic HIV infection and antigen-specific <1% it is likely that generic inflammatory stimuli are involved in the induction of TLMs irrespective of their antigen specificity.

In this study, we explored the mechanisms underlying B-cell dysfunction in chronic HIV infection, focusing on TLM B cells, which are associated with inhibitory receptor expression and impaired antigen responsiveness. Using bulk B-cell receptor (BCR) sequencing, we analyzed memory B-cell subsets in elite HIV controllers and individuals taking ART. Our findings revealed that TLMs in HIV, characterized by a CD21-CD27-phenotype, exhibit reduced mutational complexity and diversity compared to other memory B cells, suggesting an enrichment of innate-like or less mature cells within this population. A multi-omics investigation of an elite controller during an episode of transient HIV viraemia, that models early viral outgrowth such as occurs after ART interruption, uncovered dynamic immune responses. These included the emergence of non-TLM Env-reactive memory B cells, IgG targeting the HIV Env protein, and activation signatures across B-cell subsets. Single-cell analyses highlighted TLM heterogeneity, identifying two distinct subsets (TLM1 and TLM2) with varying differentiation states, IGHV4-34 usage, and mutational burdens. Additionally, pseudotime analyses suggested that TLMs comprise both innate-like and conventional memory B cells, underscoring their complexity. These findings provide novel insights into B-cell responses during HIV infection, with potential implications for vaccine and therapeutic strategies.

## RESULTS

### TLMs are significantly less diverse and mutated than other MBC subsets in PLWH

We first sought to understand if TLMs may be blocked in their ability to undergo antigen-driven evolution of their BCRs. We selected a variety of PLWH (with or without concurrent ART), including those in an elite control cohort and phenotyped their circulating B cells by flow cytometry (Fig 1A, Figure 1 supplement). There was inter-donor variation irrespective of whether the donors were in the ART or the Elite control group. However, the majority of memory B cells were consistently of a resting memory B cell phenotype and there was no obvious enrichment for a particular memory B cell subset within one group. To capture the inter-donor variability observed across both groups, we randomly selected five donors (two from the ART control group and three from the elite control group) and bulk-sorted resting (CD21^HI^CD27^HI^), activated (CD21^HI^CD27^LO^) and TLM (CD21^LO^CD27^LO^) MBCs based on their isotype (IgG versus IgM). These cells then underwent bulk BCR sequencing to assess their BCR affinity maturation, diversity and evolution. We observed that resting IgG MBCs were significantly more diverse than the other subsets, and that the diversity of IgG TLM and activated MBCs was similar (Fig. 1B). A similar trend was observed for IgM MBCs (Fig 1B), however this was not found to be statistically significant (p > 0.05) and may reflect their wider responsiveness to a range of antigens and distinct temporal germinal centre outputs.

**Figure 1:**
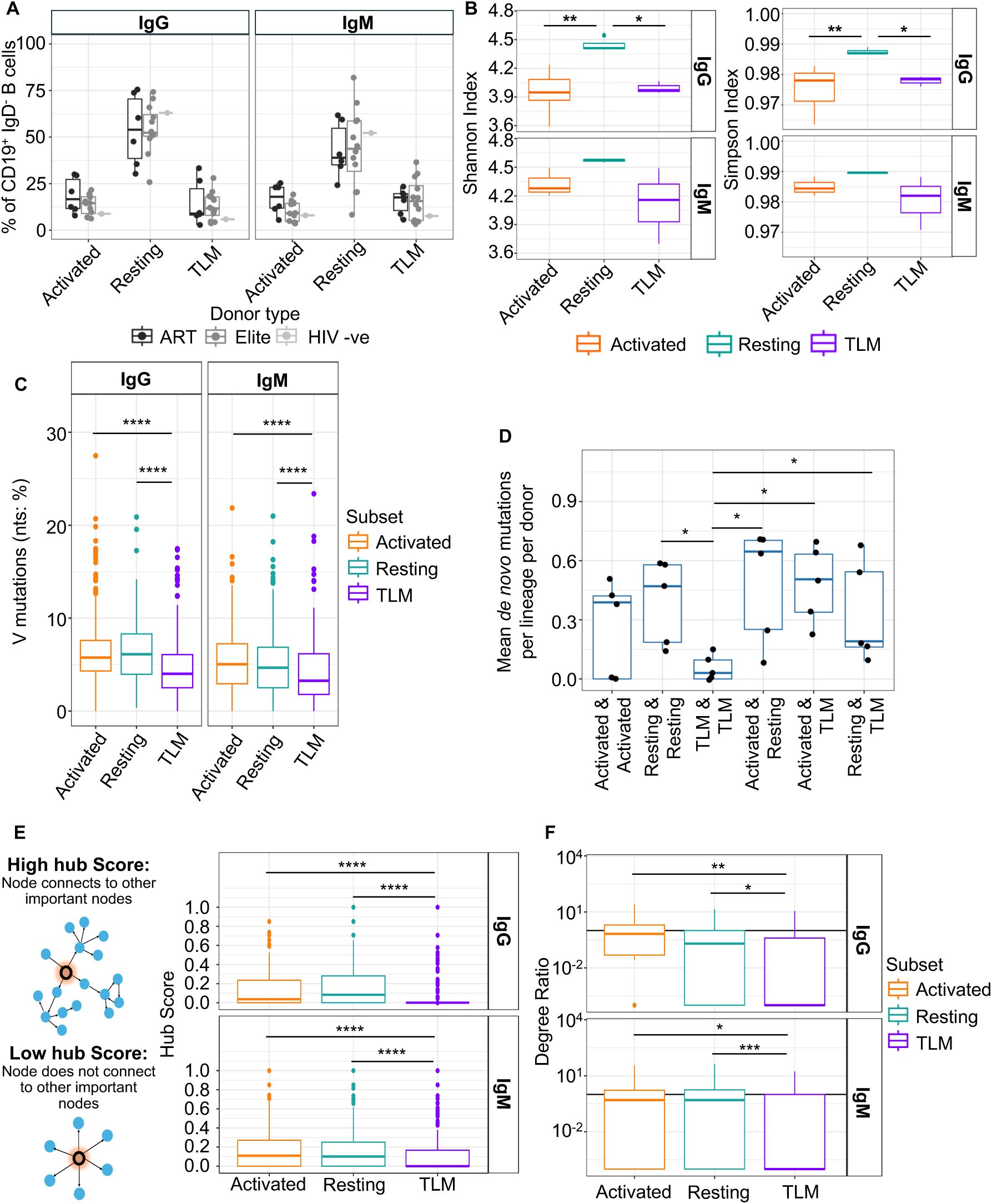
TLMs are significantly less diverse and mutated than other MBCs in HIV+ donors. (A) Flow-cytometric quantification of frequencies of IgM and IgG MBC subsets within IgD-B cells. N = 6 and 12 for ART and Elite respectively, N = 1 for HIV-ve control. (N=? per group). Gating strategy shown in Fig. S1 (B) Clonal diversity indices of MBC subsets stratified by IgG vs. IgM expression (N=5). 100 BCR sequences were randomly sampled from each donor/subset for comparison. (C) VH mutational burden (as % of nts) of BCR sequences analysed in B. (D) Calculation of de novo mutations between B cell subsets. Lineages containing multiple MBC subsets were selected, and BCR sequences underwent pairwise comparison to identify unique AA mutations (FWR1 - FWR3). The median was taken per comparison for each lineage, and then averaged to give a value per donor. The x axis shows the comparison group, and the y axis the average proportion of unique AA mutations per comparison. (E-F) BCR phylogenetic network analysis to ascertain MBC subset evolution. Networks were created for each lineage, with each node representing a BCR (self-linkages excluded). Edges between nodes were drawn if the tip-to-tip distance was below a set threshold, defined as the position of first apex in the rate of change curve for the tip-to-tip distances. The network was by only allowing edges from closer nodes to germline to further nodes from germline. Once these networks were created, the hub score (E) and out-degree to in-degree ratio (F) was calculated (hub score representation, panel E, left hand side), to provide global and local measures of node connectivity, respectively. Throughout, statistical significance was assessed using pairwise Mann-Whitney U tests with the compare_means function from the ggpubr package (p > 0.05 not marked, p < 0.05 *, p < 0.01, **, p < 0.001 ***, p < 0.0001 ****).

**Supplementary Figure 1:**
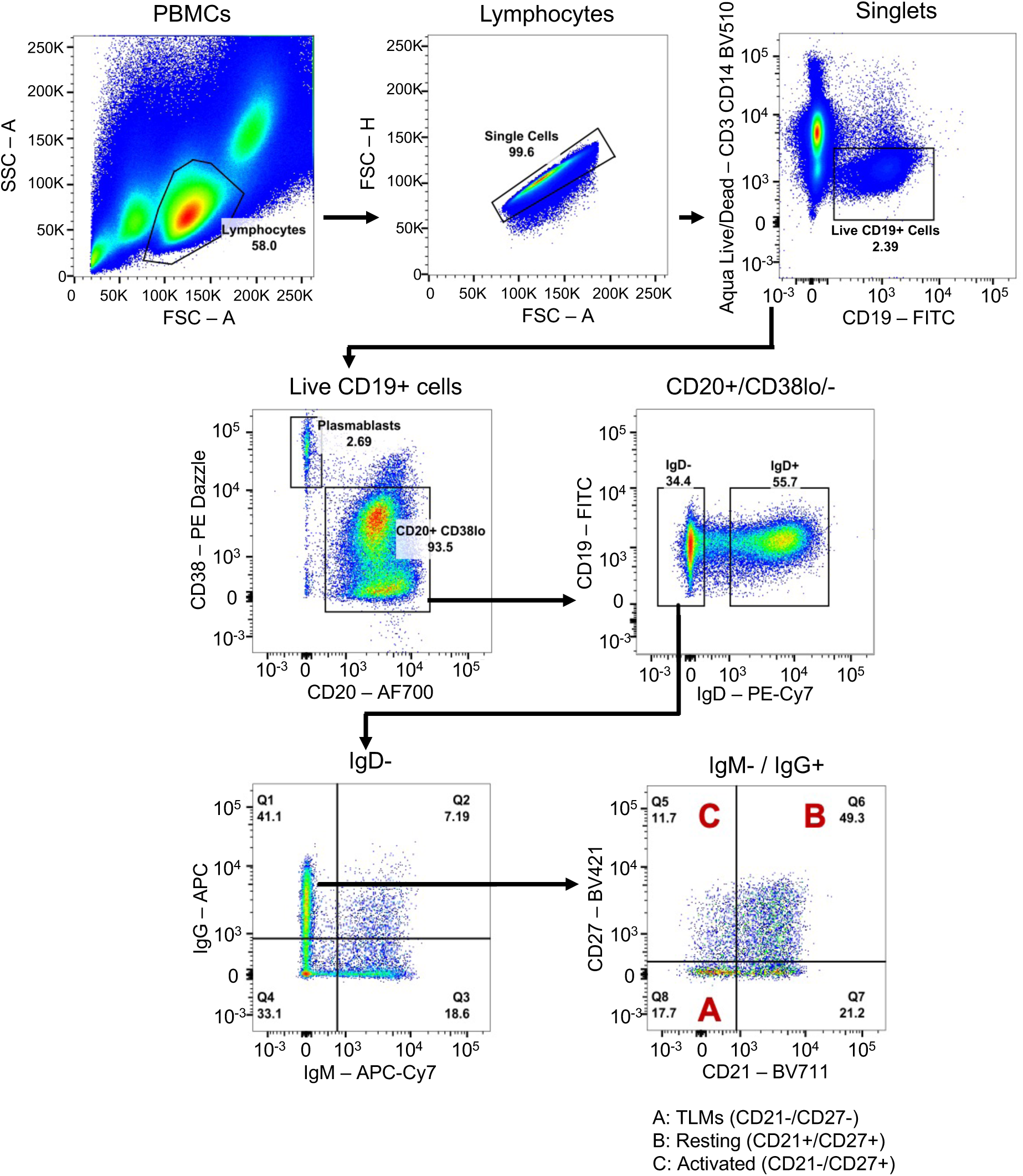
Gating strategy for the assessment and sorting of memory B cell subsets. TLM (A), Resting (B) or Activated (C) memory B cells were sorted from the IgM^+^ and IgG^+^ populations.

Next, we sought to ascertain the mutational burden of these BCR sequences across MBC subsets. Firstly, the V_H_ mutation percentage was calculated (Fig 1C). In accordance with previous work (Meffre, Louie et al. 2016), TLMs were significantly less mutated than the other MBC subsets (p = 4.86×10^-20^, p = 4.45×10^-20^, p = 8.16×10^-9^, p = 2.76×10^-4^ for IgG activated vs TLM, IgG resting vs TLM, IgM activated vs TLM, and IgM resting vs TLM respectively). This may suggest that TLMs are less able to undergo antigen-driven activation and BCR mutation. To further characterise the mutation landscape of these cells, the proportion of unique (*de novo*) AA mutations between and within each subset was compared. The median unique mutation frequency was calculated per MBC subset for each lineage, and the mean average was taken per donor (Fig 1D). This data illustrates that mutation sharing is significantly greater between TLMs than between other subsets and suggests that they are less able to undergo continued BCR diversification than other MBC subsets since they contain fewer unique mutations overall.

Next, directed phylogenetic networks were created to better understand the evolutionary trajectories of the different MBC subsets. These networks allow evaluation of evolutionary importance of each node (cell) using the hub score (Fig 1E) and the out-degree to in-degree ratio (degree ratio; Fig 1F). The hub score estimates the value of each node’s connections to other nodes within the network, i.e., the score is higher if a cell connects to another cell that is highly connected. We observed significantly lower hub scores for TLMs than other subsets, regardless of isotype (IgG; Activated p = 5.5×10^-7^; Resting p = 7.1×10^-5^, IgM; Activated p = 2.3×10^-7^, Resting p = 4.4×10^-6^, Fig 1E). This illustrates that TLM are less able to undergo continued BCR evolution than other subsets. Similarly, the degree ratios (ratio of outgoing connections divided by incoming connections) were significantly lower for TLMs across both isotypes than the other subsets (Fig 1F, IgG; Activated p = 0.0011; Resting p = 0.0308, IgM; Activated p = 0.0147, Resting p = 0.0005). Taken together, TLMs contained significantly fewer mutations, were less diverse, and were less well connected in each B cell network. Such data suggests that TLMs could be subject to additional activation/maturation controls as reported previously (Ambegaonkar, Kwak et al. 2020), a precursor population to other subsets, or produced via extrafollicular responses that undergo weaker affinity maturation (Elsner and Shlomchik 2020).

### Minor transient spike in HIV viraemia triggers a potent antibody response with expanded neutralisation breadth

Our BCR analysis of MBCs from PLWH showed that the TLM subset may have impaired capacity for BCR mutation and diversification. To address how and when TLMs arise, we focussed further study on one donor where elite HIV control was interrupted by a transient viral load and where longitudinal samples (pre-, during and post viral blip) were available. Access to such timepoints allowed us to gain a unique insight into B cell dynamics during a relatively acute and low viral blip as may be seen during ART cessation. This ART naïve individual (referred to as “index participant”) had a low/undetectable viral load (<50 copies/mL) at the time of recruitment to the elite controller cohort (month 0 – Pre Blip) in October 2013. However, when sampled at month 41 a viral load of 603 copies/mL was detected (Blip). This viral blip was associated with a CD4^+^ T cell count decrease from 950 cells/μL to 650 cells/μL (Fig 2A). Following the viral blip CD4^+^ T cell counts remained largely stable through to month 65, although they never fully recovered to the pre-blip levels. Similarly, the viral load returned to low levels of ∼100 copies/mL at month 52 (Post Blip).

**Figure 2:**
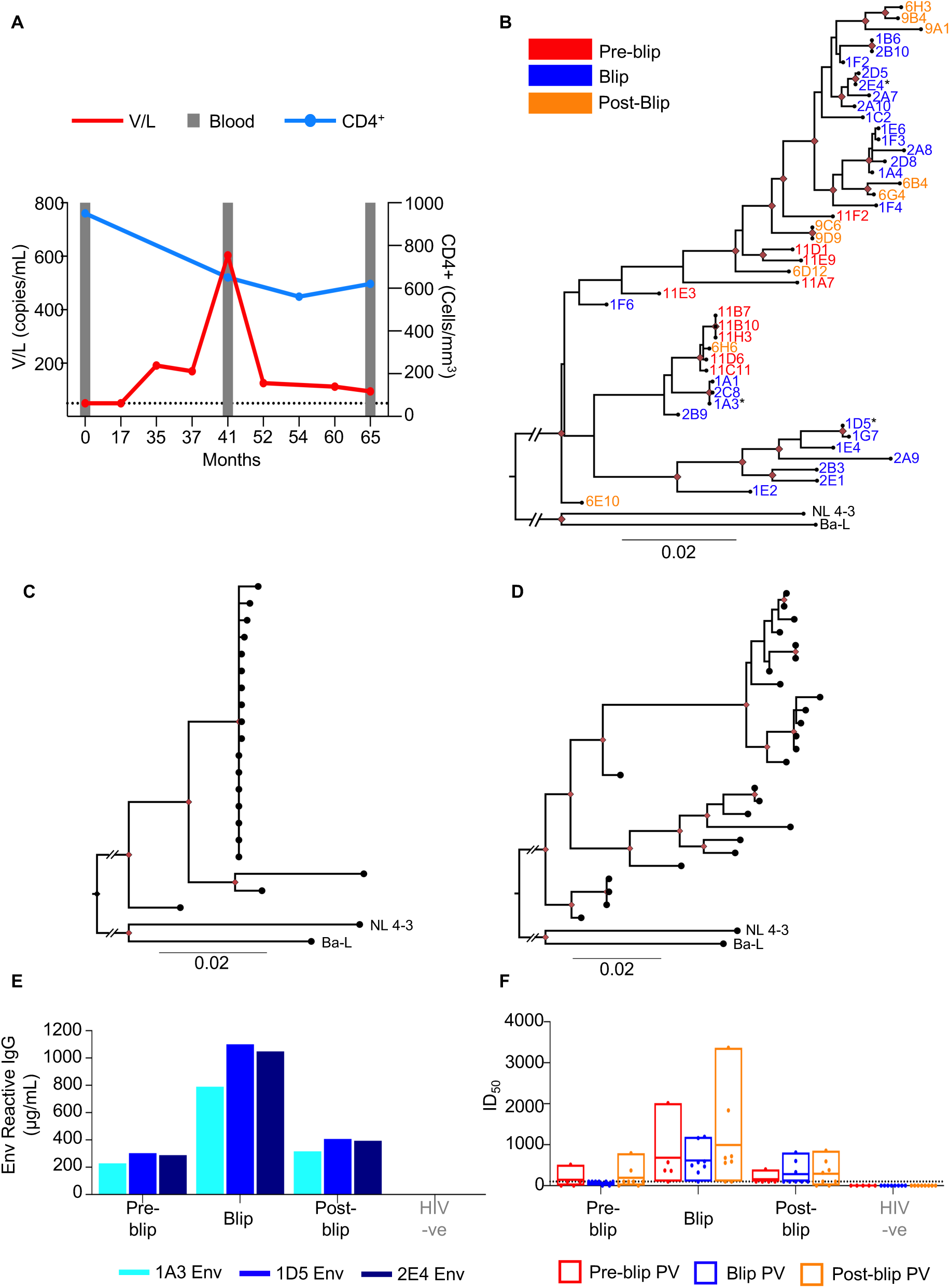
Minor transient spike in HIV viraemia triggers antibody response. (A) Dynamics of viral load (V/L, copies/mL) and CD4^+^ T cell count (per mm^3^) from an elite controller (index participant). Grey bars indicate timepoints of stored blood (PBMC/plasm) samples. (B) Maximum Likelihood Phylogenetic analysis of *env* nucleotide sequences (amplicons spanning AA35-683, HXB2 numbering) isolated from the index participant. Sequences are coloured according to the sampling timepoint. Red nodes represent bootstrap support > 70%. Clade B sequences (NL4-3 and Ba-L) were included as outgroups for phylogeny rooting. Root branches have been shortened to prevent large branches caused by differences between participant sequences and outgroups. (C-D) Maximum Likelihood Phylogenetic analysis of *env* nucleotide sequences (amplicons spanning AA35-683, HXB2 numbering) isolated from an individual on ART at one timepoint (C) or the index participant at the viral blip timepoint (D). Red nodes represent bootstrap support > 70%. (E) Anti-Env IgG titre in the plasma from the index and a HIV-negative participant. Plasma titre was assessed via semi-quantitative ELISA against the 1A3, 1D5 and 2E4 Env proteins (encoded by *env* sequences isolated from the index participant). (F) Pseudovirus neutralisation 50% inhibitory titres (ID_50_) of plasma from the index and HIV-negative participant, respectively. Pseudovirus was expressed using autologous *env* sequences cloned from the index participant including 6 pseudoviruses from the pre blip timepoint (month 0), 8 pseudoviruses from the blip timepoint (month 41) and 8 psuedoviruses from the post blip timepoint (month 65). The dotted line represents the detection limit of the assay (ID_50_ =1:100). If no neutralisation was detected, samples were assigned an ID_50_ of 0. If only a single data point of the plasma dilution series was above 1:100, no accurate ID_50_ estimation could be determined and thus samples were assigned an ID_50_ of 1:100. The box plot lines show the median and 10-90^th^ percentile.

**Supplementary Figure 2:**
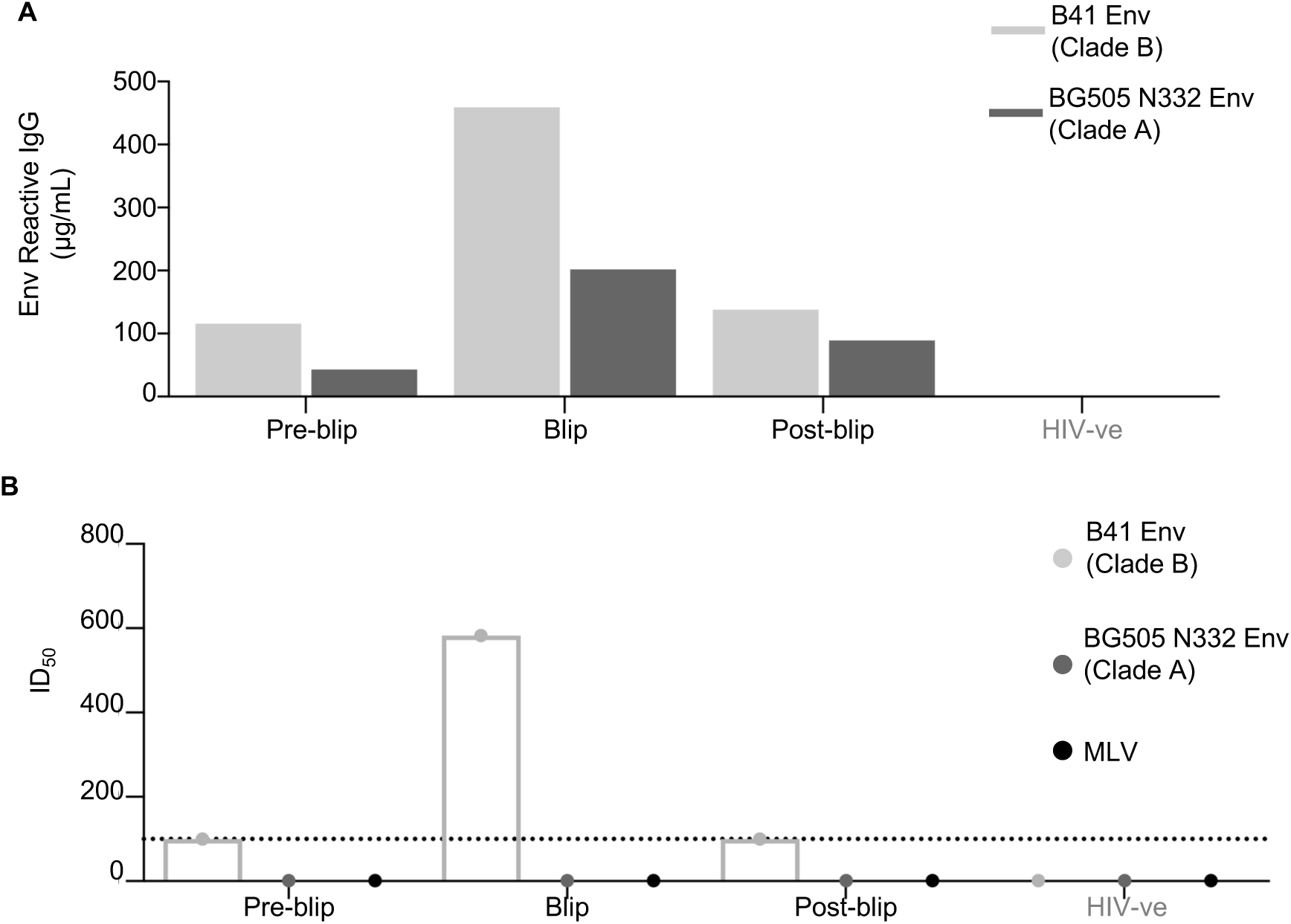
(A) Anti-Env IgG plasma titre of the index and a HIV negative participant assessed against the heterologous B41 Env (Clade B) and BG505 N332 Env (Clade A). (B) Pseudovrius neutralisaton 50% inhibitory titres (ID_50_) of plasma from the index and a HIV negative participant against the heterologous B41 (clade B) and BG505 N332 (clade A) *env* sequences. MLV encoding pseudovirus was included as a control to confirm the absence of ART.

Analysis of the integrated HIV *envelope (env)* gene sequences by single genome amplification (SGA) from genomic DNA revealed a high degree of sequence variation within the index participant across pre-blip, blip and post-blip timepoints (Fig 2B). While certain branches of the phylogenetic tree were enriched with sequences from specific time points, the sequences from the post-blip time point were dispersed throughout the tree, suggesting limited selective pressure across the assessed timepoints. Interestingly, in contrast to diverse *env* sequences from the viral blip (index participant) (Fig 2D), as anticipated sequences recovered from a different PLWH who was on ART (Fig 2C) were largely clonal, likely driven by the proliferation of infected T cell clones rather than viral outgrowth and diversification (Bozzi, Simonetti et al. 2019). Of note, the sequence diversity of the index participant viral quasi-species during the Blip may have been underestimated because of the limited RNA recovery at low viral load (<1,000 copies/mL).

To profile the B cell response to the viral blip we first determined the anti-Env plasma IgG titre. Env is the only viral antigen exposed on the surface of the HIV virion and is thus the major target for the B-cell mediated antibody response, including neutralising antibodies against HIV that are the goal of many HIV vaccine candidates. Three *env* sequences representative of the sequence variation and glycosylation features present in the index participant were selected, cloned and expressed as stabilised soluble recombinant proteins (1A3, 1D5 and 2E4) by utilising SOSIP mutations (Sanders, Derking et al. 2013). The stabilised Env proteins were used to quantify the autologous anti-Env IgG titre and compared with the binding response against two non-autologous Env SOSIP proteins: BG505 N332 (Clade A) and B41 (Clade B) (Sanders, Derking et al. 2013, Pugach, Ozorowski et al. 2015). Strikingly, the anti-Env IgG response of the index participant rose to >1 mg/mL at the viral blip (Fig 2E). As expected, the titre against autologous Env proteins (Fig 2E) was higher than that against heterologous proteins and no response was observed in HIV-negative control plasma (Fig 2 Supplement A). Interestingly, the IgG titre against the heterologous proteins peaked at the viral blip, suggesting that the small viral load increase was capable of boosting both the autologous and heterologous IgG titre.

Subsequently, plasma neutralisation was assessed using a pseudovirus assay as previously described (Sarzotti-Kelsoe, Bailer et al. 2014, Gupta, Abdul-Jawad et al. 2019). The inhibitory dilution 50 (ID_50_) values highlight that plasma neutralisation was broadest and most potent against autologous pseudoviruses at the viral blip (Fig 2F), in line with the ELISA readings (Fig 2E). This is in contrast to weaker and less broad responses at the pre blip and post blip timepoints (Fig 2F). Interestingly, the viral blip timepoint plasma was unique in neutralising a heterologous pseudovirus; B41, albeit at a low level (Fig 2 Supplement A). Overall, the neutralisation profile of the index participant appeared to be the weakest in terms of breadth and potency against autologous pseudoviruses prior to the viral blip (Fig 2F), followed by a ∼5-10 fold increase in potency and increased breadth associated with the viral blip, before the response appears to partially return to baseline (Fig 2F).

Taken together, these data show that in an elite controller, even a minor spike in viral load (603 copies/mL) can induce a potent but short-lived neutralising plasma antibody response against HIV. This makes the index participant a suitable case for studying B cell response to HIV viraemia, including TLM development with particular relevance in the acute setting of ART cessation.

### Transient viral blip induces expansion of Env-reactive IgG^+^ memory B cells

After characterising the humoral response to this minor viral blip, we next examined potential changes in the MBC compartment. We used fluorochrome-tagged autologous 1A3 and 1D5 recombinant Env proteins to detect antigen-specific memory B cell at the blip and post-blip timepoints. Unfortunately, limited PBMC availability precluded such analysis in the pre-blip sample. Env^+^ IgG+ MBCs formed almost 1% of MBCs at the blip timepoint (Fig 3A-B). However, this population proportion contracted by 75% at the post-blip timepoint as the index participant regained viral control (Fig. 3B), mirroring our ELISA and neutralisation data (Fig. 2E-F). Surprisingly, the flow-cytometric phenotyping of all IgG^+^ memory and Env^+^ IgG^+^ B cells at both timepoints demonstrated a marked similarity to an HIV-negative donor MBC phenotype and not a donor with chronic viraemia (Fig 3C), with resting subset representing >75% and TLM < 10% of MBCs, respectively (Fig. 3C). Although there was no enrichment of TLM cells in the Env^+^ subset as has been previously reported for PLWH with high chronic viraemia (Meffre, Louie et al. 2016), we did observe a more than doubling of the activated Env^+^ subset proportion compared to activated cells in the Env^-^ subset (Moir, Malaspina et al. 2008). As expected, the donor with chronically high HIV viraemia had a clear expansion of TLMs (Fig 3C).

**Figure 3:**
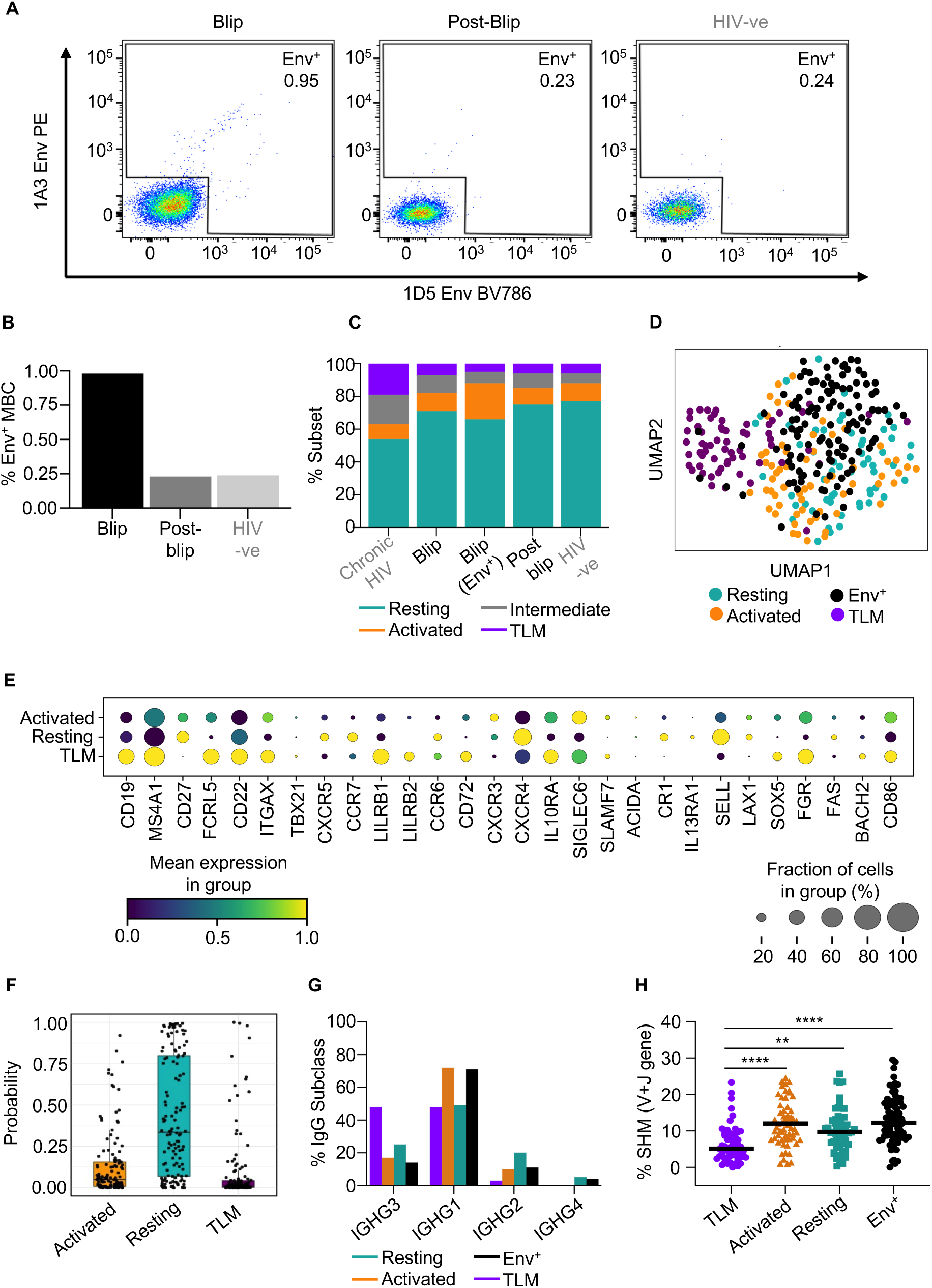
Transient viral blip induces expansion of Env-reactive IgG^+^ memory B cells. (A) Gating strategy for FACS-sorting of 1A3 Env-PE and 1D5 Env-BV786 specific memory B cells (Aqua live/Dead^-^, CD4^-^, CD19^+^, IgM^-^ IgG^+^) from index participant at the blip (month 41) and post-blip (month 65) timepoints, or a HIV negative participant. (B) Percentage of Env-reactive B cells within IgG^+^ MBCs stratified by participant. (C) Proportion of MBC subsets (resting, activated, TLM, intermediate) within total IgG^+^ memory B cells across three donors (Chronically viraemic, elite controller and HIV-negative) and timepoints (elite controller only: Blip and post-blip). The analysis was based on flow-cytometric phenotyping of the expression of CD21 and CD27. Proportion of Env+ B cell subsets is calculated within total Env+ IgG+ B cells. (D) UMAP visualization of single cell transcriptomes from TLM, activated and resting MBCs (n=301) sorted from the elite controller post blip and Env+ memory B cells from the blip timepoint. Single cell libraries were prepared via the Smart-seq 2 pipeline (Picelli 2019), a total of 88 resting, 88 activated, 88 TLM and 161 Env^+^ MBC are profiled. Cells are annotated by their sorting identity. (E) Expression of selected genes associated with memory B cell subsets profiled in D. The fraction of cells is shown by the dot size and the mean expression level is reflected by the colour. (F) Similarity of single-cell transcriptomes of Env^+^ IgG^+^ memory B cells with the resting, activated and TLM IgG^+^B cell subsets sequenced in D. Similarity is calculated as probability using the Celltypist annotation tool trained on the sorted post-blip MBC subsets. (G-H) IgG subclass usage (G) and nucleotide somatic hypermutation levels (V+J gene) (H) of sequenced memory B cell populations shown in D. One way-ANOVA with Tukey’s multiple comparison used for statistical testing. *p>0.032, **p>0.021, ***p>0.0002 and ****p>0.0001.

Next, we used FACS to isolate IgG+ cells from all three memory subsets (resting, activated and TLM) and also Env-reactive IgG+ MBCs for single-cell RNA/BCR sequencing to assess their transcriptomic similarity. UMAP clustering showed TLM as a distinct population, whereas there was a significant overlap between the resting and activated MBC subsets (Fig 3D). Interestingly, the Env reactive cells largely clustered with the resting and activated subsets but not TLMs (Fig 3D), mirroring their flow-cytometry phenotype (Fig 3C). TLM population identity/annotation was confirmed by their expression of bona fide TLM marker genes such as FCRL5 and ITGAX and low expression of CD27 when compared to the resting and activated subsets (Fig 3E). We subsequently used the automated annotation tool Celltypist to quantify the transcriptional similarity of the Env-reactive cells to MBC subsets. This confirmed that the vast majority of Env-reactive cells were transcriptionally most similar to resting memory B cells, with only a small proportion of cells resembling TLMs (Fig 3F).

Finally, we compared the differences in the use of IgG subclasses and levels of SHM among MBCs. In line with our earlier data (Fig 1C), TLM cells have significantly lower levels of SHM than resting and activated memory B cells (Fig 3H). TLM cells also displayed decreased usage of downstream IgG subclasses such as IgG2 and IgG4, suggesting a less mature or more innate-like phenotype (Fig 3G). Again, Env-reactive MBCs had a BCR profile (SHM, IgG subclasses) more similar to resting/activated memory B cells (Fig 3G, H).

Overall, these data confirm that the viral blip was accompanied by an expansion of activated Env-reactive IgG memory B cells. However, this discrete viral blip was not large enough to induce the global memory B cell dysfunction typically associated with chronic HIV viraemia nor dysfunction within Env-reactive memory B cells. Instead, the antigen-specific memory B cells observed (presumably precursors of anti-Env antibody-secreting cells) resembled resting/activated memory B cells both phenotypically and transcriptionally. Although TLM cells were present, their frequency was significantly lower than in chronically infected individuals and more in keeping with healthy (uninfected) donors.

### Transient viral blip induces a complex response across both B- and non-B cell immune subsets

To examine the transcriptomic changes in the whole B-cell and non-B-cell immune compartment of the elite controller across all three time points in an unbiased approach, we performed single-cell RNA/BCR sequencing of MACS-selected circulating B cells spiked with all PBMCs at 3:1 ratio (favouring B cells). This generated total 48,602 single-cell transcriptomes (post-QC) with 13 distinct cellular clusters identified (Fig. 4A and C, Supplemental Fig. 4A-C and E). CellTypist automated cell annotation (Fig 4D, supplemental Fig. 4D) (Stoeckius et al., 2017) was validated and further refined using a panel of eight CITE-seq markers (Supp Fig 4. H).

**Figure 4.**
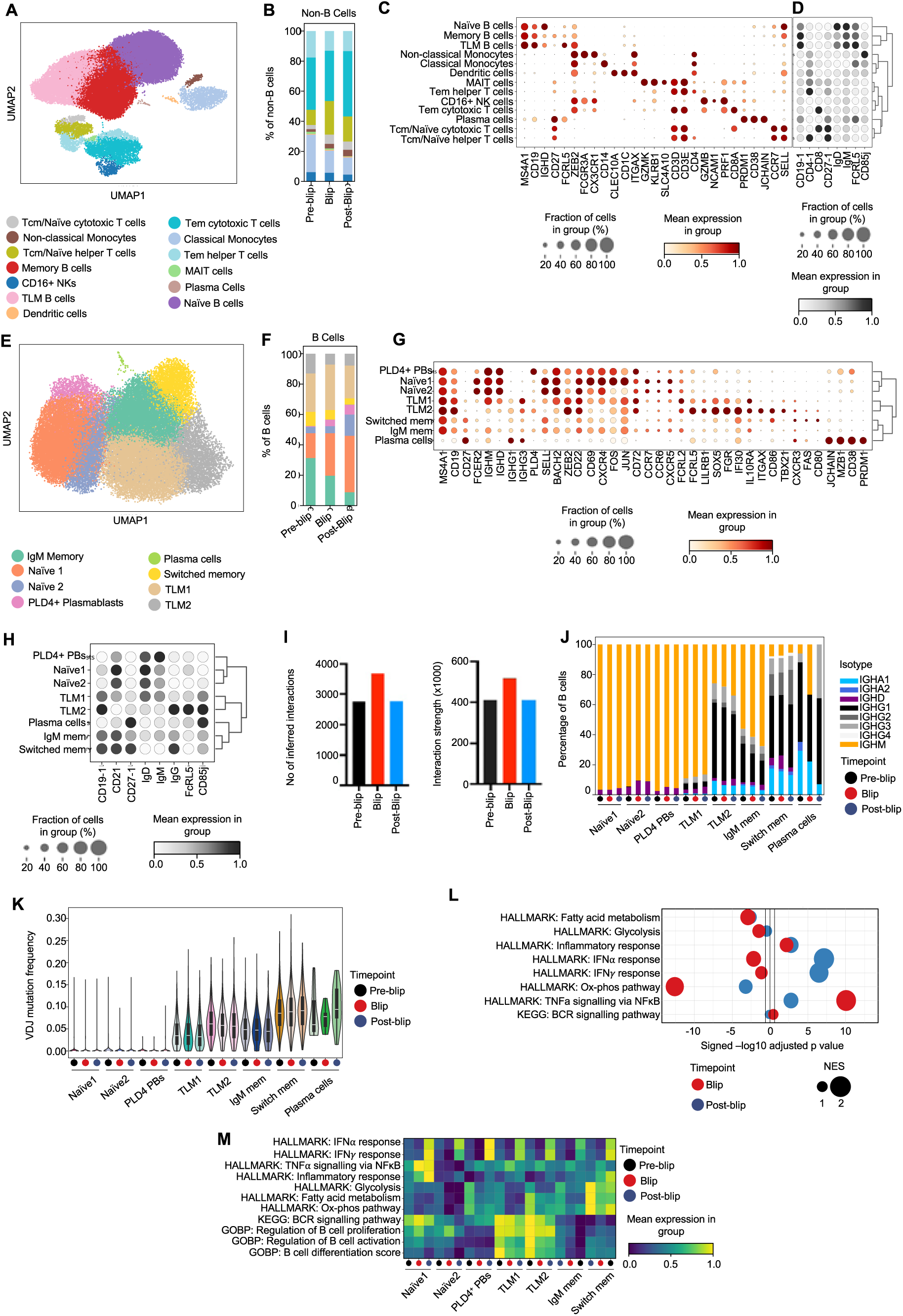
Transient viral blip induces a complex response across both B- and non-B cell subsets. (A) UMAP visualization of annotated single-cell transcriptomes (48,602 cells) recovered from circulating MACS-selected B cells spiked with PBMC in 3:1 ratio of the index participant (elite controller) across all timepoints. (B) Bar plot showing proportion (% of non-B cells) of non-B cell subsets across all three timepoints. (C-D) Dot plot showing expression of selected marker genes (C) and CITE-seq protein markers (D) across annotated cell clusters from A. The fraction of cells expressing a particular marker is shown by the dot size and the marker mean expression level is reflected by the colour. (E) UMAP visualization of finely annotated B-cell transcriptomes (42,149 cells) extracted from the dataset shown A. (F) Bar plot showing proportion (% of B cells) of annotated B cell subsets across all three timepoints. (G-H) Dot plot showing expression of selected marker genes (G) and CITE-seq protein markers (H) across annotated B-cell clusters from E. The fraction of cells expressing a particular marker is shown by the dot size and the marker mean expression level is reflected by the colour. (I) Box plots showing number of inferred cell-cell interactions (left) and interaction strength (right) across timepoints using CellChat package. Both B- and non-B cells included. (J) Isotype usage analysis depicted as % of B cells within each subset and timepoint. (K) Violin plot showing VDJ mutation frequency stratified by subset and timepoint. (L) GSEA of selected Hallmark and KEGG gene sets based on pre-ranked DEG comparing all B cells from blip (red dots) and post-blip timepoint (blue dots) to the reference (pre-blip B cells). Dot size reflects normalized enrichment score (NES). Vertical black lines indicate the threshold for statistical significance. (M) Heatmap showing scanpy expression score of selected HALLMARK, KEGG and GOBP gene sets in each B cell subset and timepoints. Scaled by row.

**Supplementary Figure 4.**
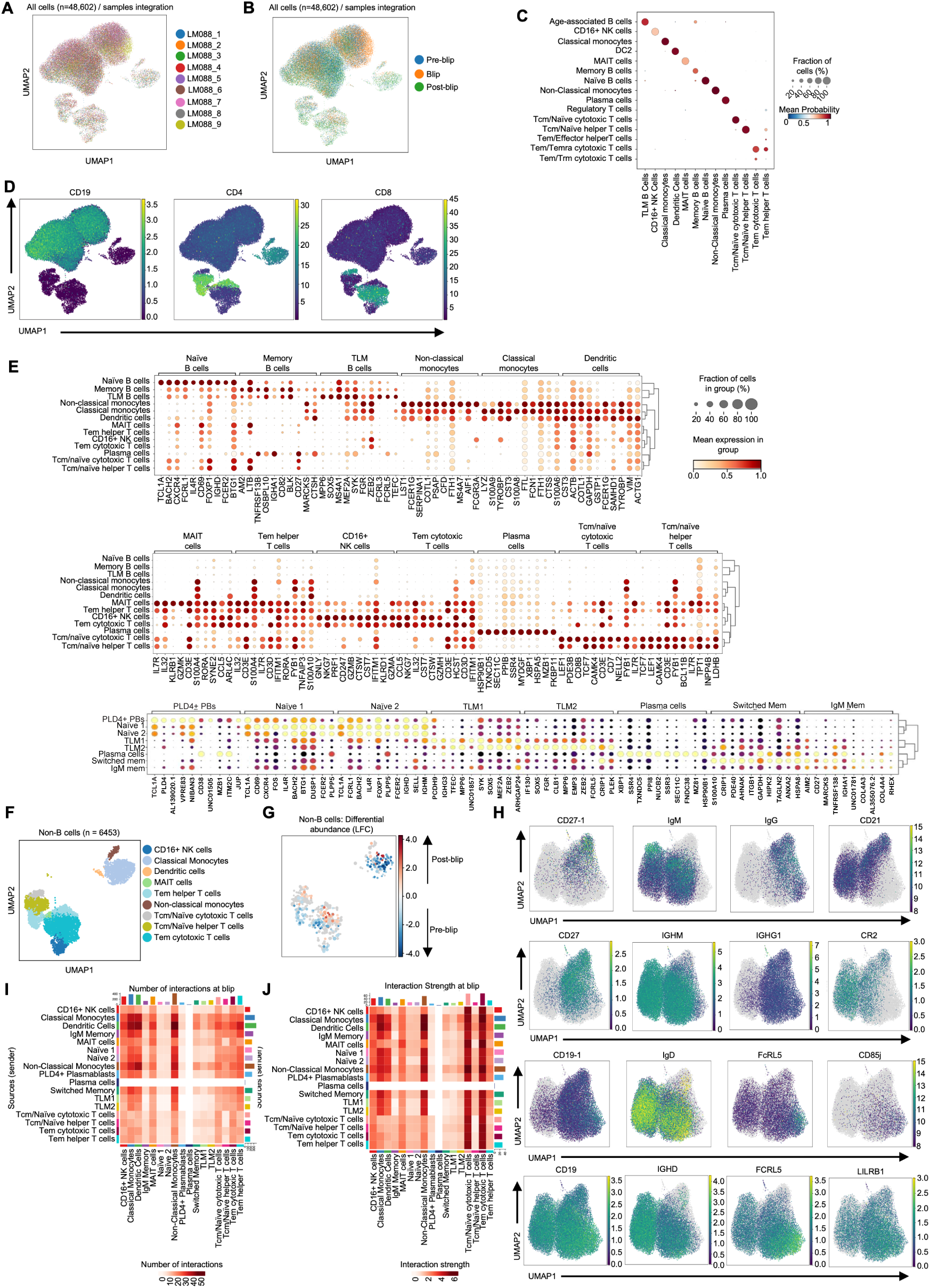
Transient viral blip induces a complex response across both B- and non-B cell subsets. (A-B) UMAP visualization of all single-cell transcriptomes (48,602 cells) as described in Fig. 4A and annotated by sequencing sample (A) and timepoint (B). (C) Comparison CellTypist automated annotation (left labels) and manually-adjusted annotation (bottom labels) of clusters identified in Fig. 4A, expressed as mean probability (circle colour). Circle size represents what fraction of manually-annotated cluster was assigned to each CellTypist-annotated clusters. (D) UMAP visualization of all cells described in Fig. 4A but coloured by their normalized expression of *CD19*, *CD4* and *CD8*. (E) Expression of top 10 marker genes for each cell subset annotated in Fig. 4A. Wilcoxon rank-sum test with Benjamini-Hochberg correction used for statistical testing, only genes with adjusted p-value < 0.05 and log-fold change >1 shown. The fraction of cells in each group expressing an indicated gene is reflected by the dot size and the mean gene expression by the dot colour. (F-G) UMAP visualization of non-B cells selected from the dataset described in Fig. 4A coloured by cell subset annotation (F) and log-fold change (LFC) in their differential abundance across timepoints (generated by milopy package). (H) UMAP visualization of B cells from Fig. 4E, comparing expression of selected markers at RNA (GEX, top plots) and surface protein level (CITE-seq, bottom plots). (I-J) Heatmaps depicting number (I) and strength (J) of inferred cell-cell interactions at viral blip across all annotated cell clusters described in Fig. 4A.

The analysis of the non-B cell compartment revealed an expansion of helper T cells during the viral blip (Fig. 4B). This was unexpected as increased viral load typically correlates with decreased helper T cells due to their direct destruction by the HIV-1 virus (McCune 2001). However, it is plausible that the relatively minor viral blip was sufficient to stimulate helper T cells expansion without causing their widespread depletion. Interestingly, in the post-blip phase, we observed a marked expansion of cytotoxic T cells (Fig. 4B) that likely played a dominant role in regaining viral control.

In the B cell compartment, we annotated eight distinct cell clusters (Fig. 4E) based on their marker genes (Fig. 4H) and CITE-seq panel expression (Fig. 4G): Naive1, Naive2, PLD4+ plasmablasts, TLM1, TLM2, IgM memory, switched memory, and plasma cells. Notably, we identified two distinct TLM populations, both expressing canonical TLM markers although with a different intensity. TLM2 cells had higher expression of ITGAX, TBX21 (T-bet), LILRB1, and CD86 (Fig. 4G). A similar expression pattern was also observed at the surface protein level by CITE-seq (Fig. 4H). This suggests that TLM1 may be a precursor to the TLM2 population, which resembles previously described TLM populations more closely (Moir, Ho et al. 2008, Holla, Dizon et al. 2021).

The analysis of the changes in the B-cell compartment across timepoints revealed an increase in naïve B cells and PLD4+ plasmablasts during blip and their persistence post blip (Fig. 4F). The emergence of PLD4+ plasmablasts may potentially explain the neutralisation observed at the viral blip timepoint. The expansion of naïve B cells during and after the viral blip may result from their enhanced generation and/or recruitment to combat the viral blip. In line with our previous flow cytometry and Smart-Seq2 data (Fig. 3), the proportion of TLM cells remained stable during the viral blip but contracted post-blip. However, within TLMs, TLM1 clearly expanded during the blip (Fig. 4F), suggesting their active role in responding to the viral blip and supporting the idea that TLM1 and TLM2 cells may have different roles.

The active coordination of antiviral immune response during the blip was further evidenced by the increased total number and strength of predicted cell-cell interactions accounting for all annotated immune cells (Fig. 4I, Fig. S4J-K). Of note, PLD4+ plasmablasts had one of the lowest number of predicted interactions at the blip timepoint, which may, together with predominant IGHM expression (Fig. 4J) and lower levels of SMH (Fig. 4K), indicate their extrafollicular differentiation pathway, allowing for a rapid anti-viral IgM response.

Similarly, TLM1 cells predominantly expressed IGHM and had lower SHM burden compared to TLM2 that were mostly switched to IgG subclasses, indicating that TLM1 cells are less mature than TLM2 cells (Fig. 4J-K). However, TLM2 were in turn less mature than switched memory as evidenced by more restricted SHM. During blip and post-blip, TLMs (unlike switched memory subset) increased their proportion of unswitched cells with a small drop in median SHM, consistent with an influx of new (less mature) B cells in response to viraemia (Fig. 4J-K).

Analysis of Hallmark gene signatures across the whole B-cell compartment across timepoints revealed a significantly enriched TNF-α signature during the blip, which declined post-blip, when an enhanced interferon-α and -γ signature emerged instead (Fig. 4L). This temporal signature pattern was seen across all B cell subsets, although with variable intensity (Fig. 4M): Naïve1 subset displayed the strongest TNF-α and inflammatory response during blip, while PLD4+ plasmablasts were the top expressor of both interferon signatures (Fig. 4M). Unexpectedly, this global blip-induced interferon response was not accompanied by the usually observed TLM expansion, for which persistent viraemia might be required. Surprisingly, TLM1 and TLM2 were the most active B-cell subset in terms of BCR signalling, activation and differentiation with only slight decrease of these signatures blip and post-blip phases (Fig. 4M). This persistent TLM activation state might indicate evolving early dysfunction, ultimately leading to reduced responsiveness, as documented in TLMs from chronically viraemic donors (Moir, Ho et al. 2008, Kardava, Sohn et al. 2018, Ambegaonkar, Kwak et al. 2020).

Taking together, our comprehensive unbiased single-cell data analysis elucidated dynamic changes in B-cell and non-B cell compartments in response to a transient viral blip. This viral blip induced TNF-α and subsequent interferon response in all B cell subsets, with naïve B cells and newly emerged (likely extra-follicular) PLD4+ plasmablasts being the most robust responders. Surprisingly, TLM2 cells did not expand in response to the transient viraemia but along with TLM1 cells displayed a persistently activated phenotype. Thus, this analysis emphasises the nuanced interplay between viral load and immune modulation in an elite HIV controller.

### TLM subsets are heterogenous and display innate-like B cell characteristics

To investigate the clonal relationship and potential differentiation pathway of annotated B cell subsets (in particular, TLMs) we utilised the single-cell BCR-seq data generated in the previous experiment. A principle component analysis based on VJ gene usage revealed that TLM1 cells across all three timepoints clustered together and separate from other B cell subsets (Fig. 5A-B). This was driven mainly by a high use of IGHV4-34 segment (Fig. 5C), which was also observed, albeit to a lesser extent, in the TLM2 subset (Fig. 5C). IGHV4-34 is often found in self-reactive and lymphoma B cells (Pascual, Victor et al. 1991, Grillot-Courvalin, Brouet et al. 1992, Parr, Johnson et al. 1994, Sebastian, Alcoceba et al. 2012). Surprisingly, TLM1 subset also appeared the most clonal apart from plasma cells, the analysis of which was likely less reliable given their small cell count (Fig. 5D). Overall, the high frequency of IGHV4-34 segment use, clonal appearance despite a relatively low SHM burden (Fig. 4K) indicates that TLM1 subset was highly enriched for innate-like B cells (Schickel et al., 2017).

**Figure 5.**
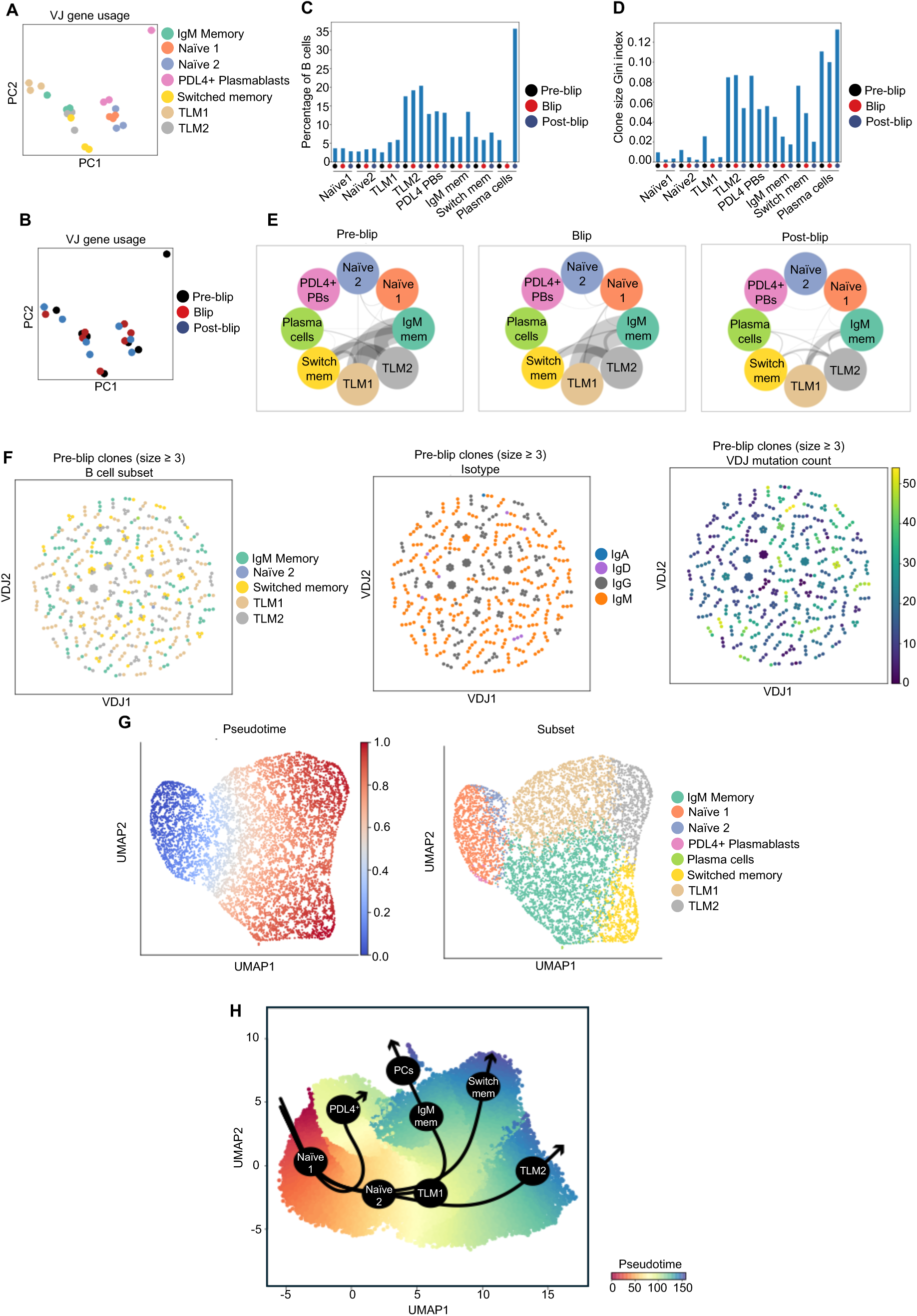
TLM subsets are heterogenous and display innate-like B cell characteristics. (A-B) PCA based on VJ gene usage by elite controller B cells coloured by B cell subset (A) and timepoint (B). (C) Bar charts showing percentage of B cells using IGHV4-34 segment within each subset and timepoint. (D) Bar charts showing clonality (clone size gini index) of each B cell subset stratified by timepoint. (E) Circos-style plots depicting clonal overlap between each B cell subset pair stratified within each timepoint. The thickness of the grey connecting line between circles reflects the number of overlapping BCR clonotypes. (F) Single-cell BCR network plots of all pre-blip clones consisting of at least three cells, coloured by B cell subset annotation (left), heavy chain isotype class use (middle) and mutation count (right). Each circle/node represents a single B cell with a corresponding set of BCR(s). Cells belonging to the same clone are connected with a black line. (G) UMAP visualisation of all B cells described in Fig. 4E, coloured by GEX-based pseudotime and overlayed by inferred developmental lineages (black arrows), generated by slingshot package. (H) UMAP generated by the BCR-seq-based pseudobulk VDJ trajectory analysis (dandelion package) of B cell dataset described in Fig. 4, coloured by calculated pseudotime and B-cell subset annotation.

The clonal overlap analysis showed the strongest link between TLM1 and IgM memory B cells, reaching its maximum during blip (Fig. 5E). Interestingly, TLM2 cells overlapped equally with TLM1, IgM memory and switched memory subsets, suggesting a likely heterogeneous origin of these cells (Fig. 5E). Indeed, we observed many clones composed of cells from different B cell subsets, although individual clones seemed mostly homogenous in their isotype use (Fig. 5F). TLM2 cells within the clones had the highest VDJ mutation count, suggesting them as the differentiation endpoint (Fig. 5F).

To explore the differentiation pathways of the B cell subsets further, we used the Slingshot package to infer pseudotime and developmental trajectories (Fig 5H). We identified four distinct developmental lineages (Fig. 5H). PDL4+ plasmablasts differentiated directly from naïve B cells, which is consistent with their proposed extrafollicular origin. Interestingly, the TLM1 subset appeared to be the branching point for three district lineages (IgM memory, switched memory and TLM2). Although TLM1 formed a cluster of transcriptomically similar B cells, they are still heterogenous and poised for different differentiation pathways. These cells likely include innate-like B cells that tend to progress into TLM2, and naïve B cells differentiating into memory cells (either IgM or switched). Although switched memory B cells and TLM2 appeared as two distinct differentiation endpoints, we demonstrated earlier (Fig. 5F) that switched memory B cells can still progress into TLM2, making TLM2 the ultimate endpoint. Finally, we replicated these GEX-based pseudotime analysis findings also using BCR-seq-based pseudobulk VDJ trajectory analysis (Fig. 5G).

In conclusion, our investigation into the origins and differentiation potential of TLM cells through BCR and 5’GEX data analysis revealed high IGHV4-34 usage, high clonality with lower SHM burden compared to memory B cells, suggesting their particular enrichment for innate-like B cells. However, both TLM subsets also contained cells from IgM memory and switched memory clones, indicating the potential for TLM1 cells to differentiate into conventional memory B cells whilst TLM2 appeared to be the final differentiation endpoint.

## DISCUSSION

Our study provides crucial insights into B cell function during HIV infection, with a particular focus on the TLMs that have been implicated in HIV-associated B-cell dysfunction. Through bulk BCR sequencing of circulating B cells from multiple PLWH, we consistently observed that TLMs exhibited significantly lower SHM burden but also lower diversity compared to other MBC subsets, indicating either an early differentiation state, a blockage in antigen-driven evolution and/or innate-like phenotype. Hence, while some TLMs may serve as precursors for more mature B cell subsets, some of these cells may be in a terminal differentiation state. We used multi-omics approaches to investigate B-cell/TLM response to HIV viraemia in a unique case of an elite controller who experienced a transient HIV viral blip and show that the TLM population is a heterogenous population with some exhibiting innate-like B cell features, some appear to be a precursor population for activated and resting memory B cells whilst some appear to be a differentiation endpoint.

We observed an increase in anti-Env IgG and associated expansion of Env-reactive IgG+ memory B cells without significant changes in the TLM abundance during the viral blip. The Env-reactive memory B cells displayed a predominantly non-TLM phenotype, clustering transcriptionally with resting and activated MBCs. This was in contrast to previous studies of chronic HIV viraemia, where Env-reactive cells were enriched for TLMs and total TLMs were suggested to contribute to overall immune dysfunction (Moir, Malaspina et al. 2001, Moir, Ho et al. 2008, Moir, Buckner et al. 2010). The minimal expansion of TLMs during the blip suggests that short-lived increases in viral load may not drive the generation of dysfunctional TLM as seen in chronic infection but instead support the production of functional, antigen-specific memory B cells. This finding is in line with the recent study by Cooper et al. that showed that during acute LCMV infection in mice there are more antigen-specific memory B cells and fever CD11c^+^Tbet^+^ DN B cells than in chronic LCMV infection (Cooper, Xu et al. 2024). They also found that prolonged exposure to high levels of type I interferon associated with viraemia induced memory B cell dysfunction via epigenetic modifications, which were reversible only in the first few week post infection (Cooper, Xu et al. 2024).

However, TLMs were present by classical cell surface expression markers and using sc-RNA sequencing, we identified two distinct TLM subpopulations: TLM1 and TLM2. The former were predominantly IgM-biased, less mutated and expressed inhibitory receptors at a lower level than TLM2. Pseudotime analysis suggested that TLM1 is likely a precursor population, with the potential to progress into more mature B cell states (either classical memory or TLM2). In contrast, TLM2 cells, which were predominantly IgG+, appeared to represent a terminal differentiation state. The TLM2 gene expression pattern closely resembled the expanded TLM population described previously during chronic HIV infection (Moir, Ho et al. 2008, Holla, Dizon et al. 2021) and the double-negative (DN2) population found in healthy controls (Stewart, Ng et al. 2021). In both previous studies, these subsets were proposed to be the differentiation endpoints in agreement with our findings. Although this study was just in one individual the identification of multiple TLM subsets is consistent with previous observations using scRNA sequencing technologies (Stewart, Ng et al. 2021).

One intriguing observation from our study was the strikingly high use of IGHV4-34 in TLM1 and to a lesser degree in TLM2 population, which is often associated with self-reactive and innate-like B cell phenotypes (Schickel, Glauzy et al. 2017, Ray and Rothstein 2023). Innate-like B cells are heterogenous unconventional B cells characterised by their innate sensing, T-independent responding capacity and spontaneous production of germ-line encoded natural antibodies that play a role in first-line defence and tissue homeostasis (e.g., apoptotic cell disposal). Natural antibodies are typically of IgM isotype, self-reactive and have low mutational burden (Gronwall and Silverman 2014). The prime examples of innate-like B cells are marginal-zone B cells and B-1 cells, the existence of which in humans remains controversial (Suchanek and Clatworthy 2023). Innate-like B cells express negative BCR co-receptors to allow their unique tonic BCR signalling and activation pattern. Indeed, our Hallmark GSEA analysis revealed that, unlike other subsets, TLMs were in a state of persistent activation and BCR signalling across all three timepoints (highest in the pre-blip phase) suggesting an innate-like character. Hence, it is possible that this ongoing activation might be a physiological feature of innate-like TLMs rather than a sign of evolving dysfunction in this particular case which may differ in chronic infection. However, our study clearly demonstrated that TLMs contained not only innate-like B cells but also cells from the conventional memory B cell lineages, further highlighting the complexity of the whole TLM compartment.

Another B cell subset with innate-like features identified by this study were PLD4+ plasmablasts, which significantly emerged with the viral blip likely via extrafollicular route (Tull, Pitcher et al. 2021, Siu, Pitcher et al. 2022, Al-Aubodah, Aoudjit et al. 2023). PLD4 is Phospholipase D4 which is an RNA/DNA exonuclease located in endo-lysosomes of plasmocytoid dendritic cells but can be upregulated in B cells upon TLR7/9 signalling (Yasaka, Yamazaki et al. 2023). Recently, self-reactive PLD4+ plasmablast enrichment was found in patients with SLE and were shown to phenotypically overlap with T-bet+ DN2 B cells (Yasaka, Yamazaki et al. 2023). Although this subset had the strongest expression of the type I interferon signature in the post blip phase and likely produced first-line IgM, its significance for regaining viraemia control and its potential for differentiation into TLM remains to be explored during chronic HIV infection.

The primary limitation of our study is that our detailed analysis was conducted in a single elite controller experiencing a transient viral blip, which limits the generalizability of our findings. However, the temporal samples from this individual did allow us to investigate the effects of an acute and small HIV viral load increase, as may happen during treatment interruption, and its impact on humoral immunity. Furthermore, while we observed key insights into B cell dynamics, the absence of sustained/increasing viraemia in our index participant prevents us from fully exploring the mechanisms driving TLM expansion and dysfunction seen in chronic untreated HIV infection. Future studies involving larger cohorts and longitudinal data will be crucial to validate these findings and explore the pathways governing TLM development under different viral load conditions. Additionally, our analysis of the viral sequences was constrained by inability to obtain viral RNA during the blip, limiting the investigation of the viral dynamics driving the humoral response. Nevertheless, our use of gDNA was supported by multiple previous studies suggesting that provirus from gDNA does reflect recent viral integration sequences (Brodin, Zanini et al. 2016, Bruner, Murray et al. 2016, Abrahams, Joseph et al. 2019).

Overall, our study demonstrates that TLMs generated in the context of HIV infection are a heterogeneous population significantly enriched for B cells with innate-like characteristics that are interferon not antigen driven. Two distinct TLM subsets (TLM1 and TLM2) can be distinguished: TLM1 (T-bet^low^) cells retain the capacity for further differentiation, whereas TLM2 (T-bet^hi^) cells appear to represent a terminal differentiation state. Our data suggest that such small transient increase in HIV viral load, while capable of eliciting a functional immune response (including PLD4+ plasmablasts), does not drive the expansion of dysfunctional TLMs typically associated with chronic HIV infection which potentially supports on going clinical trials using brief therapy interuptions. Understanding the tipping point between when viraemia promotes functional immune responses versus inducing the emergence of dysfunctional B cell subsets like TLMs and enhanced T cell destruction is essential for developing strategies to enhance vaccine efficacy and therapeutic interventions in chronic HIV infection. Future research should aim to elucidate the molecular signals and pathways that govern B cell differentiation and dysfunction in the context of both acute and chronic HIV infection.

## AUTHOR CONTRIBUTIONS

Conceptualization: L.E.M.

Investigation and Data Analysis: L.M., O.S., P.T., S.A.G., C.R-S., E.T., C.P., Z.K.T. and R.P.

Single-Cell and bulk RNA Sequencing Data Analysis: L.M., O.S., P.T. and Z.K.T.

Patient Recruitment: R.K.G.

Writing – Original Draft: L.M., O.S. and L.E.M.

Writing – Review and Editing: L.M., S.A.G., O.S., P.T., C.R-S., E.T., M.Y, C.P., Z.K.T., R.P., M.Z.N, K.J.D., R.K.G., M.R.C. and L.E.M.

Funding Acquisition: L.E.M.

## ACKNOWLEDGEMENTS

LEM is supported by a Medical Research Council Career Development Award (MR/R008698/1).

## MATERIALS AND METHODS

### Ethics/Patient information

The index participant was classified as exhibiting elite control based upon their largely stable low viral load (<100 copies/mL) over 13 years whilst not on ART (patient preference) prior to the viral blip. Samples from PLWH who had detectable viraemia or non-ART mediated viral control were collected as part of a protocol approved by local research ethic committee (London – City & East REC 12/LO/1572). Samples from PLWH on ART (suppressed) and HIV-negative participant samples were previously collected and processed with ethical approval by South Central – Hampshire B (REC 19/SC/0423)(Touizer, Alrubayyi et al. 2023). Studies complied with all relevant ethical regulations for work with human participants and conformed to the Helsinki declaration principles and Good Clinical Practice (GCP) guidelines. All subjects enrolled into the studies provided written informed consent.

### PBMC Isolation

Heparinised blood was isolated, diluted with an equal volume of PBS (Sigma Aldrich, Dorset, UK) and layered over 20 mL Histopaque-1077 (Sigma Aldrich). Peripheral blood mononuclear cells (PBMCs) and plasma were then isolated by centrifugation (400x g, 20 minutes, no brake). A 1x 15 mL aliquot and 2x 1 mL aliquots of plasma were recovered and stored at −80°C for future use. The mononuclear layer was then recovered, washed in 40 mL PBS and collected by centrifugation (400x g, 5 minutes). Cells were then washed in complete RPMI (RPMI-1640 media (Sigma Aldrich) supplemented with L-glutamine (Sigma Aldrich), penicillin/streptomycin (Sigma Aldrich) and 10 % fetal bovine serum (FBS) (Sigma Aldrich)), pelleted by centrifugation (400x g, 5 minutes), resuspended in complete RPMI and counted using trypan blue exclusion. PBMC were cryopreserved in cryovials for future use by resuspension at 1×10^7^ cells/ml in FBS 10% DMSO (Sigma Aldrich) and stored at -80°C.

### Single Genome Amplification (SGA) of *Envelope* genes

The Qiagen DNEasy kit (Qiagen, Manhester, UK) was used to extract DNA from 1×10^7^ PBMCs using the spin column protocol with some minor modifications. Cells were lysed for 30 minutes at 56 °C rather than the recommended 10 minutes. A longer lysis step was found to lead to increased DNA yield. Additionally a modification was made to the elution step, whereby 90μl of 5mM Tris-HCL (Sigma Aldrich) was dispensed onto the column, the column was then warmed to 40 °C and left for 20 minutes prior to centrifugation at 13,000 RPM for 1 minute. HIV *envelope* single genomes were PCR amplified as previously described [17] using isolated genomic DNA as template. In brief, gDNA was diluted in PBS and nested PCR carried out using High Fidelity Platinum Taq DNA polymerase (Invitrogen, Paisley, UK) with primers Env5out and OMF19, followed by nested primers Env5in and Env3in (Carlson, Schaefer et al. 2014, Flerin, Bardhi et al. 2019). PCR amplifications were carried out in 1× High Fidelity Platinum PCR buffer, 2mM MgSO4, 0.2mM dNTPs, 0.2μM of each primer and 0.025 units/μLPlatinum Taq, with a final reaction volume of 10 μL. The thermocycler protocol was as follows: 94°C for 2 minutes, followed by 35 cycles of 94°C for 15 seconds, 55°C for 30 seconds and 68°C for 4 minutes, finishing with a 10 minute elongation step at 68°C. 2kb *env* amplicons were visualised by agarose gel electrophoresis on precast 96 well 1% agarose E-gels (Invitrogen). Wells where a single *env* amplicon was obtained then underwent one final round of PCR amplification using the Env Cloning Fwd and Env Cloning Rev primers which generate an *env* amplicon compatible with Gibson cloning into the psvIII expression vector. Conditions for the final PCR were unchanged although reactions were scaled up to 25 μL.

### Envelope cloning

*env* PCR amplicons were purified using the QIAquick PCR purification kit (Qiagen) following the manufacturer’s recommendations. psvIII vector containing the HXB2 *env* sequence was digested with the BamHI-HF and KpnI-HF restriction enzymes (NEB, Hitchin, UK) at 37°C for 3 hours in 1x CutSmart buffer (NEB). The digested plasmid was then visualised by agarose gel electrophoresis and the digested plasmid purified using the QIAquick Gel Extraction Kit (Qiagen) following the manufacturer’s recommendations. Purified inserts were then cloned into the digested psvIII expression vector via Gibson assembly. Briefly, 100 ng of digested psvIII vector was incubated with a 3 fold molar excess of PCR insert in 1x NEBuilder® HiFi DNA Assembly Master Mix (NEB) for 2 hours at 50°C. NEB® 10-beta Competent *E. coli* (High Efficiency) were subsequently transomed with the assembly mix and incubated on LB-Agar ampicillin selection plates overnight at 37°C. The following day single colonies were upscaled to 1 mL LB-Broth ampicillin culture for 6 hours at 30°C, 200 RPM. 2 μL of culture was then added to 48 μL H_2_O and placed at -80°C for 1 hour before incubation at 95°C for 15 minutes. The resulting supernatant was then used as template for a PCR reaction with the Env Cloning Fwd and Env Cloning Rev primers as described above. *env* amplicons were visualised by agarose gel electrophoresis on precast 96 well 1% agarose E-gels (Invitrogen). Wells which contained *env* amplicons were upscaled to 100 mL LB-Broth ampicillin culture 30°C, 200 RPM for 16 hours and then plasmid DNA isolated using the plasmid plus midi sample kit (Qiagen).

### Envelope Sequencing and Phylogenetic analysis

Cloned *env* amplicons underwent Sanger-sequencing using the A-K sequencing primers. Sequence reads were trimmed at the 5′ and 3′ ends until base calls consistently reached a quality score ≥30, resulting in 99.9% accuracy of base calls. Contigs were then generated using SnapGene Software (www.snapgene.com) by alignment to the HXB2 *env* sequence. *env* contigs were trimmed resulting in amplicons from amino acid 35 to amino acid 683 (HXB2 numbering). *env* sequences were then aligned at the protein sequence level using MUSCLE v.3.8.31 (Edgar 2004) and mapped back to nucleotide, with minor manual adjustment. 2 clade B sequences (NL4-3 and B41) were included for reference and 2 clade A sequences (BG505 and 92UG029.EC1) from the Los Alamos HIV Sequence Database (http://www.hiv.lanl.gov/) were included to serve as an outgroup. A maximum likelihood phylogeny was estimated using IQ-tree software (Nguyen, Schmidt et al. 2015), with the general time-reversible nucleotide substitution model and gamma-distributed rate heterogeneity. Clade support was estimated from 1,000 nonparametric bootstrap replicate datasets. The maximum likelihood phylogeny was rooted on the outgroup branch and visualized using FigTree v.1.4.4 (http://tree.bio.ed.ac.uk/software/figtree/).

### Soluble Envelope protein production

Three diverse *env* sequences from the maximum likelihood phylogeny: 1A3, 1D5 and 2E4 were selected to be expressed as soluble stabilised proteins. To allow for stabilised expression, SOSIP mutations were introduced into the sequences as described by Sanders et al (Sanders, Derking et al. 2013) and sequences ordered as synthetic Gene Fragments with Gibson cloning compatible overhangs (Genewiz, Bishop’s Stortford, UK). p818 plasmid was digested with the BstBI and PstI restriction enzymes and gel extracted, *env* Gene Fragments were then cloned into the digested vector as described in the “envelope cloning” methods section. Stablised Env proteins were expressed in HEK293F cells and purified with either PGT145 or 2G12 affinity chromatograph followed by size exclusion chromatography (SEC) using a HiLoad 16/600 Superdex pg200 (GE Healthcare) as described previously (Sanders, Derking et al. 2013, de Taeye, Ozorowski et al. 2015, Torrents de la Pena, Julien et al. 2017). The Avi-tagged proteins were biotinylated using the BirA enzyme (Avidity, Colorado, USA) according to the manufacturer’s instructions.

### Pseudovirus production

HEK-293T cells were co-transfected with Env-expressing psvIII plasmid and a HIV Env deficient backbone plasmid (pSG3△Env) at a mass ratio of 1:2 using Polyethylenimine (PEI) max (Polysciences, Bergstrasse, Germany) at a DNA:PEI mass ration of 1:3 and incubated for 48 hours at 37°C 5% CO_2_. The pseudovirus containing supernatant was then harvested and passed through a 0.45 μm syringe filter (Sigma Aldrich), aliquoted and stored at -80°C. Subsequently, frozen pseudovirus was titrated using the HeLa-TZM-bl reporter cell assay with the Bright-Glo^TM^ Luciferase assay kit (Promega, Hampshire, UK) and the 50% tissue culture infectious dose (TCID_50_/mL) calculated.

### *In vitro* Neutralisation Assays

Pseudovirus neutralisation assays were carried out using the 96-well plate TZM-bl cell-based assay described by Sarzotti-Kelsoe et al (Sarzotti-Kelsoe, Bailer et al. 2014), with Bright-Glo^TM^ luciferase reagent (Promega) and final readout analysed using a BioTek Synergy^TM^ H1 plate reader (Agilent technologies, Manchester, UK). Briefly, plasma samples were heat-inactivated by incubation at 56°C for 1 h and then titrations of plasma starting at 1:100 incubated with luciferase encoding pseudovirus (at 200 TCID_50_/mL) in duplicate for 1 hour at 37°C 5% CO_2_. 10,000 HeLa TZM-bl cells were then added to each well and incubated for 48 hours at 37°C 5% CO_2_ before supernatants were removed and luciferase readout used as an indicator of infectivity. 100% infectivity was defined by wells which contained virus and TZM-bl cells only, with no antibody present.

### Semi-quantitative Total Ig ELISA

Half-area 96-well MaxiSorp plates (VWR, Leicestershire, UK) were coated with either 25 μL goat anti-human F(ab)’2 (1:1,000) (Jackson immunoResearch, Cambridgeshire, UK) in PBS for total IgG plates, or 25 μL mouse anti-human Ig κ light chain and mouse anti-human Ig λ light chain (1:200 each) (Becton Dickinson Biosciences, Wokingham, UK) in PBS for total IgG3 plates, and incubated overnight at 4°C. Plates were washed with PBS-T and blocked with 100 μL assay buffer (PBS, 3% BSA, 0.05% Tween-20) for 1 hour at room temperature. 25 μL of serially diluted plasma (1:100 to 1:10^7^) in duplicate or known concentrations of IgG or IgG3 standard in triplicate (Sgma Aldrich) were then applied to the plates. Plates were washed with PBS-T and 25 μL of detection antibody diluted in assay buffer added to each well: goat anti-human IgG-AP (1:1,000) (Sigma Aldrich) or goat anti-human IgG3-AP (1:1,000) (Cambridge Bioscience, Cambridge, UK) and incubated for 1 hour at room temperature. Subsequently, plates were washed 6 times with PBS-T and 25 µL of colorimetric alkaline phosphatase substrate added (Sigma Aldrich). Absorbance was measured at 405 nm after 60 minutes. Total IgG or IgG3 concentrations in plasma were then calculated based on interpolation from the IgG or IgG3 standard curves using a four-parameter logistic (4PL) regression curve fitting model.

### Semi-quantitative Env ELISA

Columns 1-9 of half-area 96-well MaxiSorp plates were coated overnight at 4°C with streptavidin in PBS (2 µg/ml per well in 25 µL) (Sigma Aldrich). Columns 10-12 were coated with either 25 μL goat anti-human F(ab)’2 (1:1,000) in PBS for IgG standard curves, or 25 μL mouse anti-human Ig κ light chain and mouse anti-human Ig λ light chain (1:200 each) in PBS for IgG3 standard curves. Plates were then washed with PBS-T and blocked with 100 μL assay buffer (PBS, 3% BSA, 0.05% Tween 20) for 1 hour at room temperature. 25 μL of biotinylated Env protein (2 µg/ml) in assay buffer was then added to streptavidin coated wells and 25 µL of assay buffer added to wells in columns 10-12 for 2 hours at room temperature. Following washing 25 μL of serially diluted plasma (1:100 to 1:10^7^) in duplicate or known concentrations of IgG or IgG3 standard in triplicate were then applied to the plates. Plates were washed with PBS-T and 25 μL of detection antibody diluted in assay buffer added to each well: goat anti-human IgG-AP (1:1,000) or goat anti-human IgG3-AP (1:1,000) and incubated for 1 hour at room temperature. Subsequently, plates were washed 6 times with PBS-T and 25 µL of colorimetric alkaline phosphatase substrate added. Absorbance was measured at 405 nm after 60 minutes. Env reactive IgG or IgG3 concentrations in plasma were then calculated based on interpolation from the IgG or IgG3 standard curves using a four-parameter logistic (4PL) regression curve fitting model.

### Flow Cytometry and single cell sorting

Fluorescence-activated cell sorting (FACS) of Env reactive cryopreserved PBMCs was performed on a BD FACS Melody (BD Biosciences). Alternatively memory B cell subsets were sorted using a BD FACSAria II (BD Biosciences). To generate fluorescently tagged Env probes 2 μg of biotinylated Env protein was incubated with either 0.5 μg streptavidin-conjugated PE (BD Biosciences) or 0.5 μg streptavidin-conjugated BV786 (BD Biosciences) for 30 minutes. Cryopreserved PBMC were thawed on ice and incubated with 100 μL of Zombie Live/Dead dye (diluted 1:400 in PBS) (Biolegend, London, UK) for every 5×10^6^ cells at room temperature for 20 minutes. The dye was then quenched by addition of complete RPMI, cells collected by centrifugation (400x g, 10 minutes) and then washed once in PBS. Cells were then incubated with specific combinations of the phenotyping antibodies +/- biotinylated tetramers listed in table 2 in a final volume of 100 μL in PBS (concentrations used were for every 5×10^6^ cells). Compensation controls were prepared according to manufacturer’s instructions using Anti-Mouse Ig, κ and negative control compensation particles (BD Biosciences). For Smart-Seq 2 experiments single B cells were gated according to the following successive gates: lymphocyte, single cells, CD19^+^ Aqua live/Dead^-^ CD4^-^ and IgM^-^ IgG^+^. From this population 1x 96 well plate of each of the following subsets was sorted: CD21^-^ CD27^-^ (TLM cells), CD21^-^ CD27^+^ (activated memory cells) and CD21^+^ CD27^+^ (resting memory cells). Alternatively antigen specific cells were sorted based on positivity for either the PE-tagged or BV786-tagged Env probes. All cells were sorted using “Yield” purity mode into chilled 96-well plates (VWR) containing 4.4 μL of lysis buffer composed of oligo-dT (5′-AAGCAGTGGTATCAACGCAGAGTACT30VN-3′) (1 μM), 0.4% Triton X-100 (Sigma Aldrich), dNTPs (1mM) (Thermo Fisher, Paisley, UK), and RNAse inhibitor (4 units) (Takara Bio, London, UK). After each plate was filled, it was underwent centrifugation at 400x g for 1 minute and placed on dry ice immediately before storage at −80°C. For bulk BCR sequencing experiments single B cells were gated according to the following successive gates: lymphocyte, single cells, CD19^+^ Aqua live/Dead^-^ CD14^-^ CD3^-^, CD20^+^ CD38^lo^, IgD^-^ and either IgM^-^ IgG^+^ or IgM^+^ IgG^-^. From this population the following subsets were sorted: CD21^-^ CD27^-^ (TLM cells), CD21^-^ CD27^+^ (activated memory cells) and CD21^+^ CD27^+^ (resting memory cells). Bulk cell populations were sorted using “Yield” purity mode into chilled 1.5 mL tubes containing lysis buffer (0.3% NP40 and ∼6.67mM RNaseOUT). After sorting of each donor was complete each 1.5 mL tube underwent centrifugation at 400x g for 1 minute and was placed on dry ice immediately before storage at −80°C. Data was analyzed on FlowJo v10 (BD).

**Table 1:**
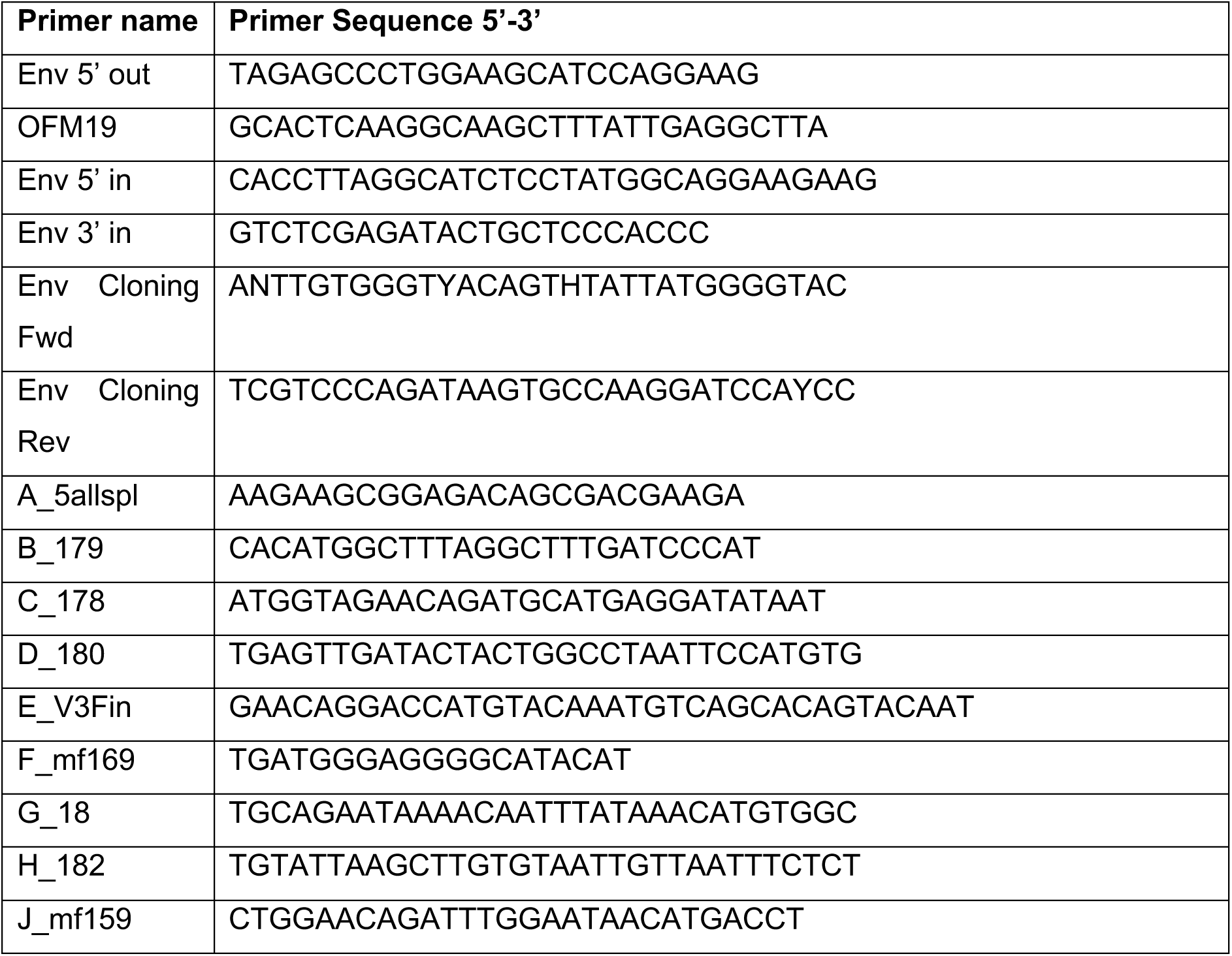
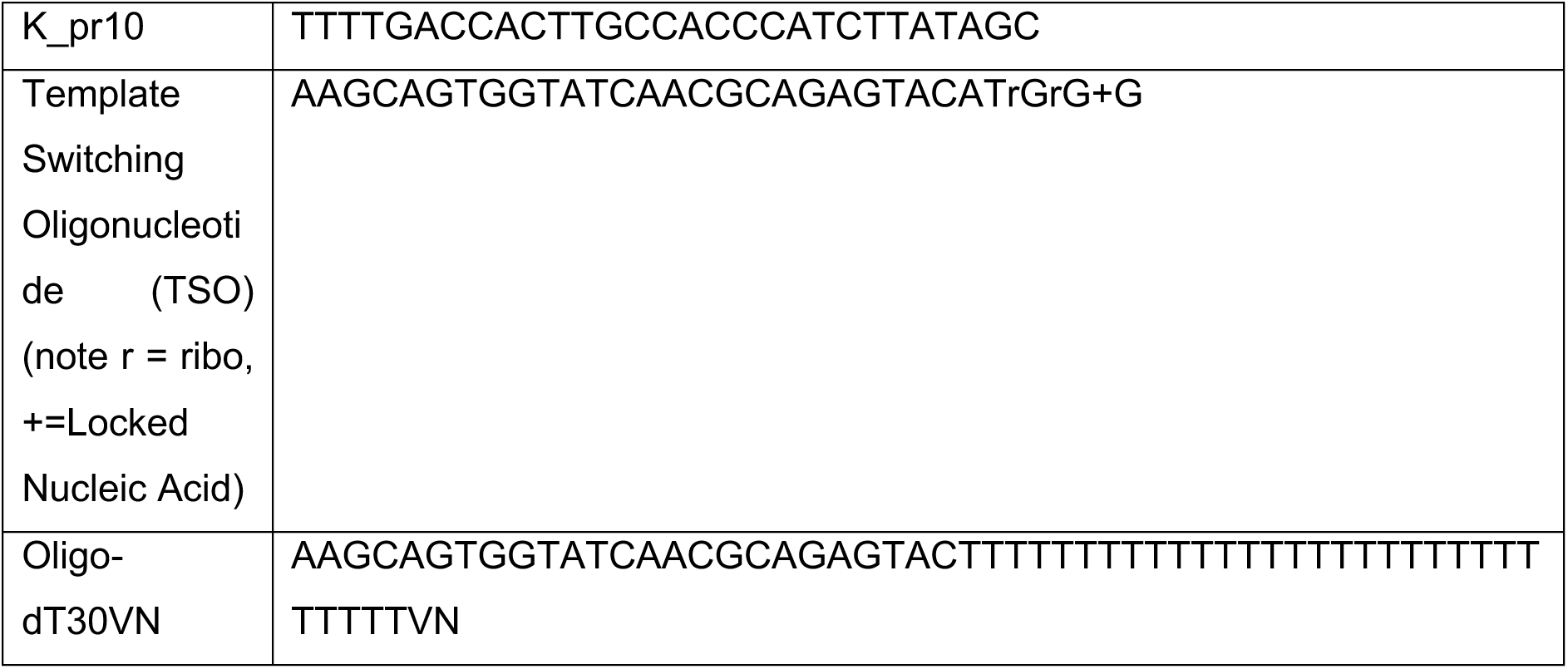
Smart Seq 2 and Env sequencing primer sequences.

**Table 2:**
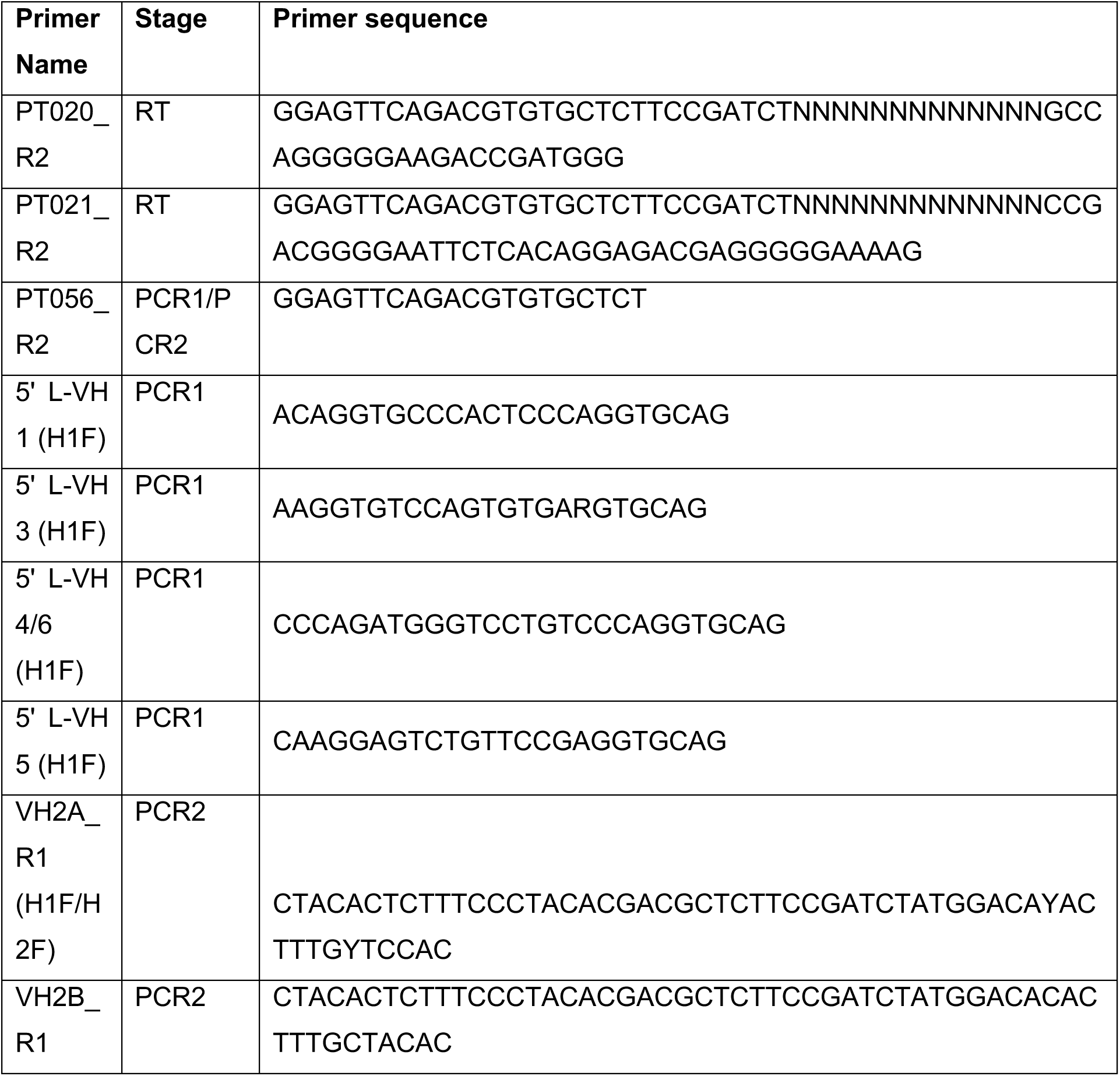
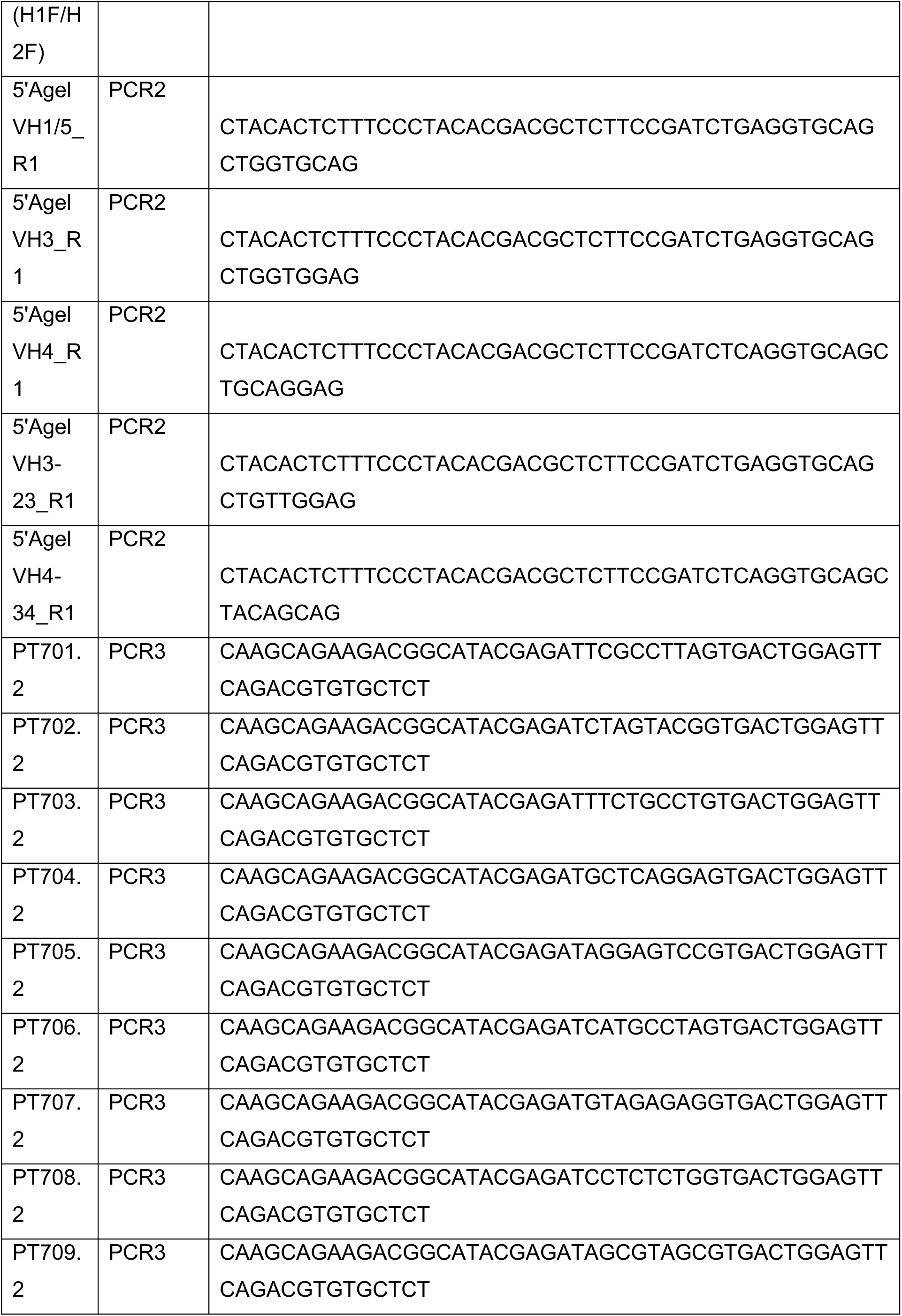
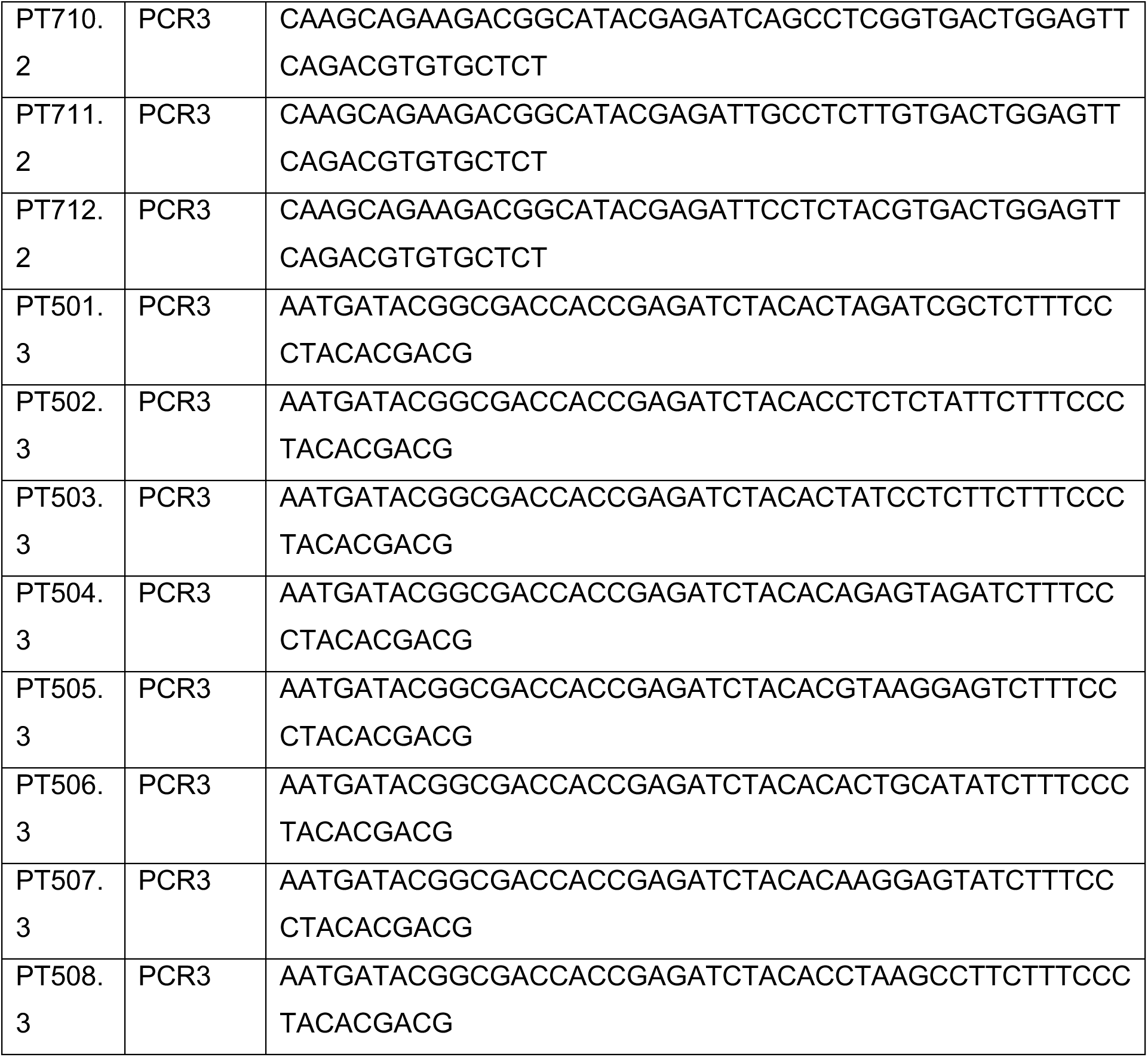
Bulk BCR Sequencing Primers.

**Table 3:**
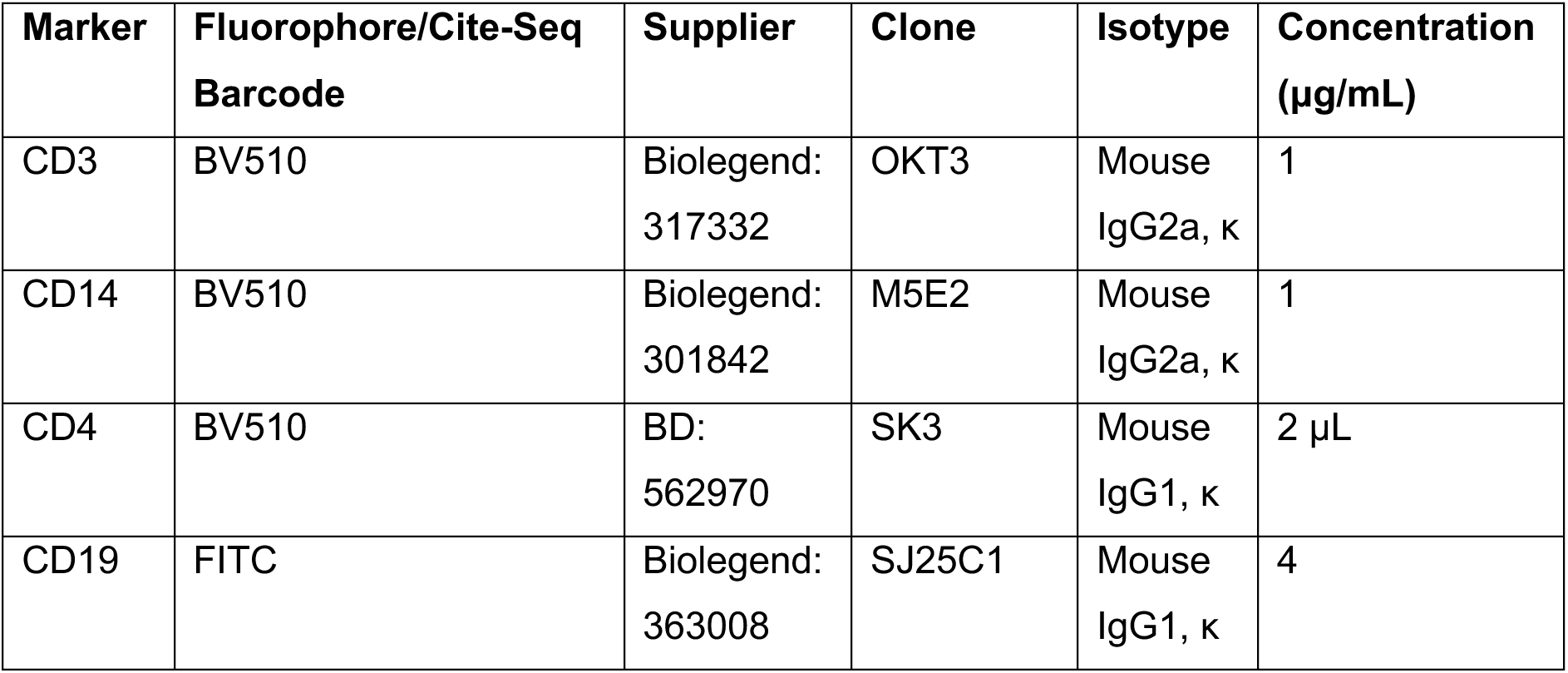
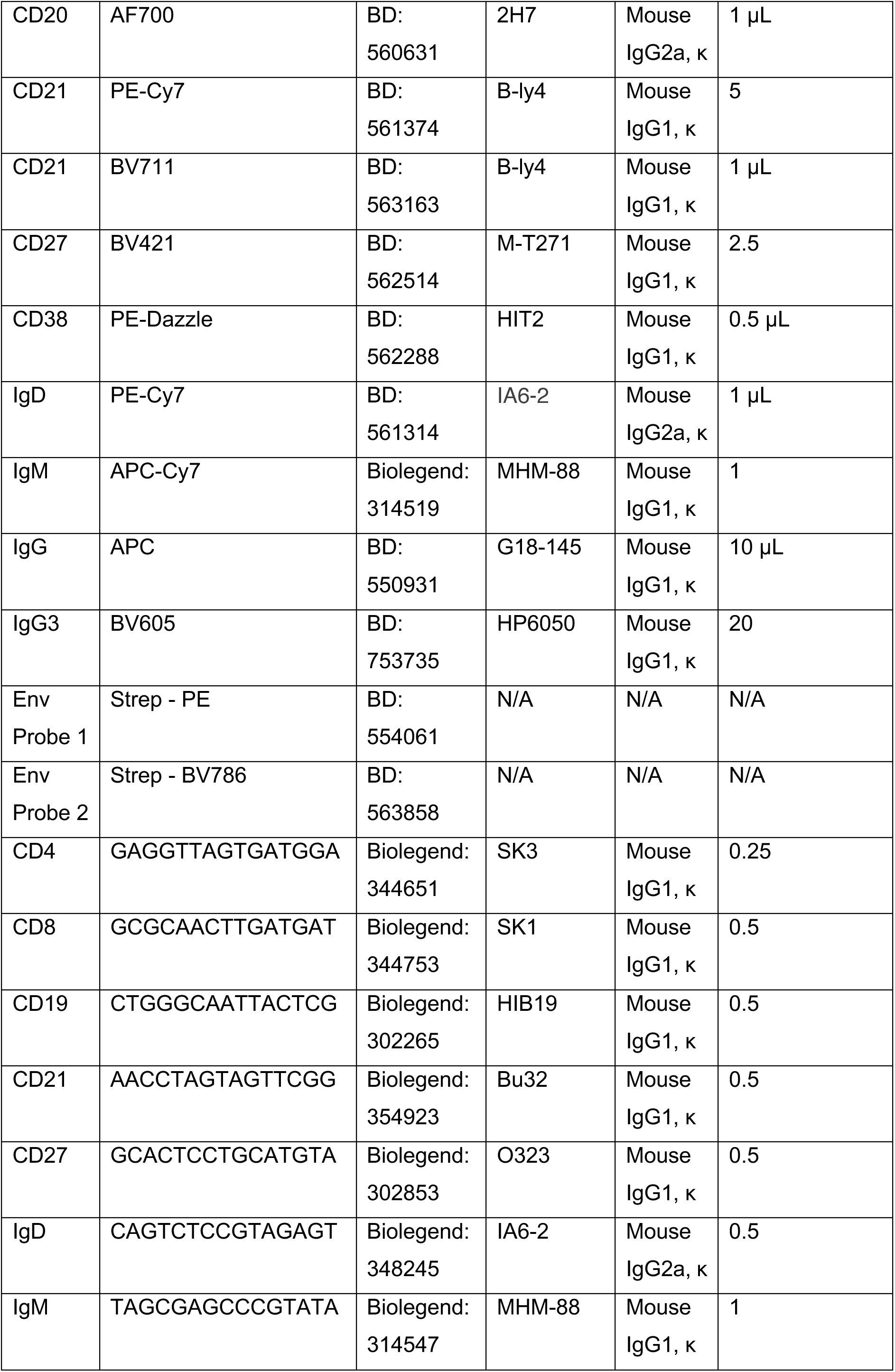
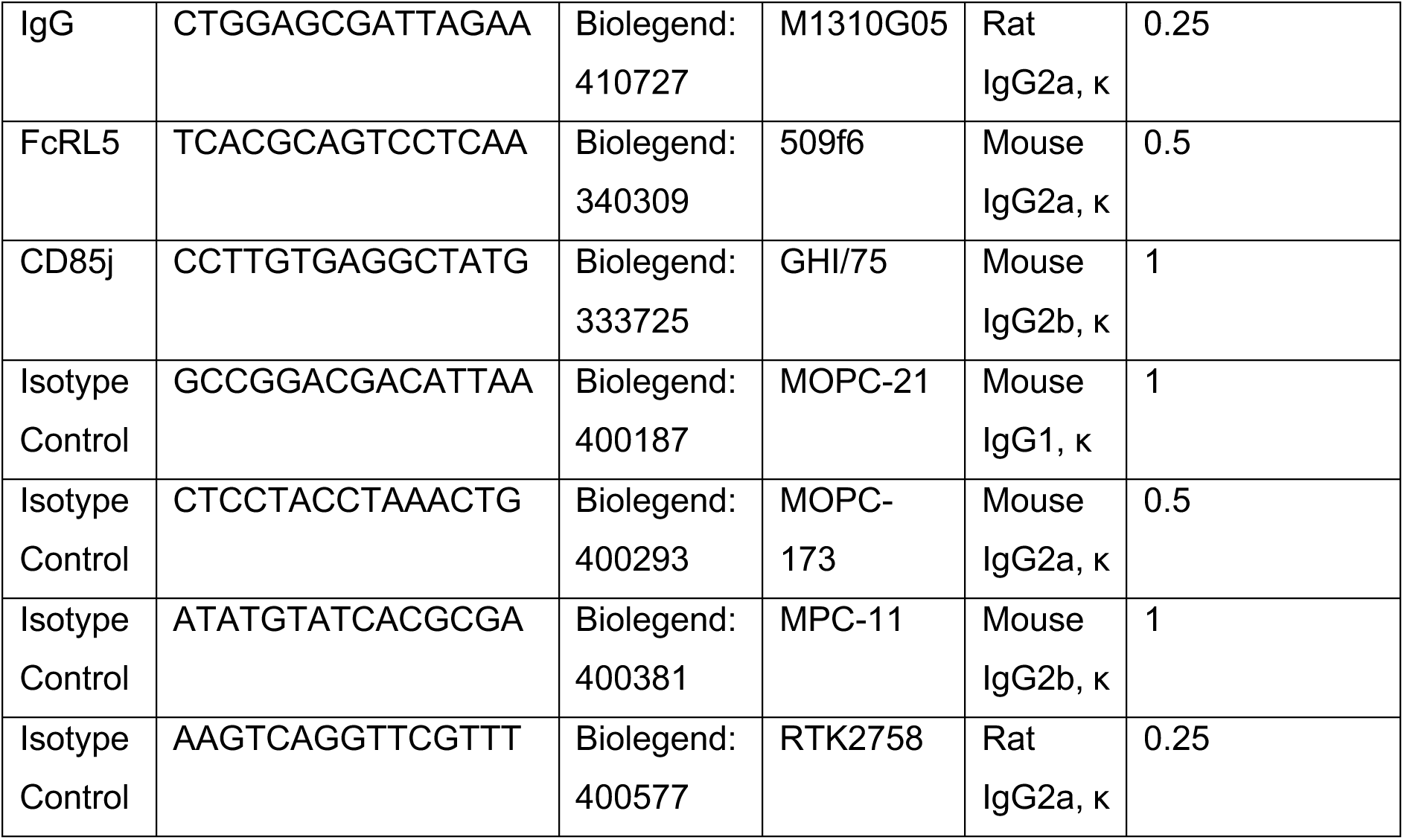
B cell phenotyping flow cytometry and CITE-Seq panels. Where suppliers do not provide antibody concentrations the volume of antibody used (μL) is provided.

### cDNA generation, library preparation, and sequencing (Smart-seq2)

The Smart-seq2 protocol (Picelli, Faridani et al. 2014) was utilised with some minor adjustments for working with lymphocytes. In brief, 96-well plates containing single cell lysates were thawed on ice. Each well then received 5.4 μL of reverse transcription mix containing Superscript II reverse transcriptase (50 units) (Thermo Fisher), RNAse inhibitor (10 units) (Tarka Bio), 5X first strand buffer (Thermo Fisher), DTT (5mM) (Thermo Fisher), betaine (1M) (Sigma), MgCl2 (12mM) (Sigma Aldrich), Nuclease-free water (Thermo Fisher) and template switching-oligo (5′-AAGCAGTGGTATCAACGCAGAGTACATrGrG+G-3′) (1μM). Reverse transcription (RT) was carried out for 90 minutes at 42°C followed by 10 cycles of: 50°C for 2 minutes and 42°C for 2 minutes to allow for completion/continuation of RT and template-switching, then finally 70°C for 15 minutes to terminate the reaction. Each well then received 15 μL of PCR mix containing Nuclease-free water, and 2x HiFi HostStart ReadyMix (Roche, Welwyn Garden City, UK). Cells were then subjected to the following thermocycler protocol: 98°C for 3 minutes, 24 cycles of: 98°C for 20 seconds, 67°C for 20 seconds, and 72°C for 6 minutes, then finally 72°C for 5 minutes. Following amplification, 0.8× AMPure XP beads (Beckman Coulter, High Wycombe, UK) were used to purify cDNA and remove primer dimers. The size distribution of cDNA in each well as well as the concentration of cDNA in the 500 to 5000 bp range was subsequently determined using a High Sensitivity D5000 ScreenTape kit (Agilent Technologies) and readout analysed on an Agilent 4200 TapeStation System. Cells that exhibited cDNA profiles indicative of degraded RNA were excluded from subsequent library preparation. The cDNA from remaining wells were diluted to approximately 0.5 ng/μL before carrying out the Nextera XT DNA library preparation (Illumina, Cambridge, UK) following manufacturer’s instructions. 0.6× AMPure XP beads (Beckman Coulter) were used to purify cDNA and remove primer dimers. Libraries were then quality checked and quantified once again using the High Sensitivity D5000 ScreenTape system and then libraries were pooled at equimolar concentrations for sequencing. In total, 443 cells were sequenced across two runs using 2×75 bp NextSeq 500 High Output v2.5 kits on an Illumina NextSeq 500.

### CITE-seq staining for single-cell proteogenomics

Frozen PBMC samples were thawed at 37 °C in a water bath. Ice cold compete RPMI was added slowly to the cells before centrifuging at 300x *g* for 5 min. This was followed by a wash in 5 ml complete RPMI. The PBMC pellet was collected, resuspend in ice cold PBS and the cell number and viability were determined using Trypan Blue. A portion of the PBMC from each timepoint was then placed in 1.5 mL tubes and placed on ice, whilst the remainng PBMC underwent magnetic cell sorting (Miltenyi Biotec, Bergisch Gladbach, Germany) to purify the B cell populations. After isolation B cells were mixed with non purified PBMC at a 3:1 ratio. The cells were then resuspend in 100 μL of mastermix containing CITE-Seq antibodies (table 2) for 30 min at 4 °C. cells were then washed three times by centrifugation at 500x *g* for 5 min at 4 °C. PBMCs were counted again and processed immediately for 10x 5′ single cell capture (Chromium Next GEM Single Cell V(D)J Reagent Kit v2 with Feature Barcoding technology for cell Surface Protein-Rev F protocol) (10x genomics, Pleasanton, CA, USA). Three lanes of 20,000 cells from each timepoint were loaded onto a 10x chip.

### Library generation and sequencing

The Chromium Next GEM Single Cell 5′ V(D)J Reagent Kit (V2 chemistry) with Feature Barcoding technology for cell surface proteins was used for single-cell RNA-seq library construction for all samples. GEX and V(D)J libraries were prepared according to the manufacturer’s protocol (10x Genomics) using individual Chromium i7 Sample Indices. The cell surface protein libraries were created according to the manufacturer’s protocol. 5’GEX, V(D)J and cell surface protein indexed libraries were pooled and sequenced on a NovaSeq 6000 S4 Flowcell (paired-end, 150 bp reads) aiming for a minimum of 50,000 paired-end reads per cell for 5’GEX libraries and 10,000 paired-end reads per cell for V(D)J and cell surface protein libraries.

### Bulk sorted BCR library construction

Bulk sorted MBC subsets (in lysis buffer) were collected from -80°C and incubated at room temperature to thaw the samples. The lysate was collected at the bottom of the tube by centrifugation. To purify the RNA, 1.7X volumes of room temperature RNAClean XP RNA purification beads were used (Beckman Coulter), binding RNA to beads for 20 minutes at room temperature. Mixtures were magnetically separated and then washed with fresh 80% ethanol. They were then air-dried before and eluted using molecular biology grade water (Ambion) for 10 minutes at room temperature. Clarified RNA containing supernatant was transferred to a clean tube.

To generate bulk BCR libraries, 15µL per RNA sample was combined with dNTPs (0.5mM, Thermo Fisher) and C-region specific primer (100nM) and molecular biology grade water, to a final volume of 17.75µL. Reverse transcription was performed using the SuperScript IV^TM^ First-Strand Synthesis System, following manufacturer’s recommendations in a final volume of 25µL, using C-region specific primers (100nM) that contained UMIs and the Illumina read 2 site. Reaction proceeded at 50°C for 45 minutes, prior to enzyme inactivation at 80°C for 10 minutes, followed by degradation of RNA:cDNA hybrids using RNaseH (New England Biolabs). cDNA product was purified using 1.0X AMPURE XP DNA purification beads (Beckman Coulter).

Eluted cDNA was supplemented with a mixture of leader sequence specific forward primers (H1F mix; 500nM), and a tag-specific reverse primer (PT056; 500nM), and 2X Q5 DNA polymerase Mastermix (New England Biolabs), to a final volume of 50µL, before undergoing PCR. This, and all subsequent PCRs, began with an initial denaturation at 98°C (50 seconds), and temperature then cycled between 98°C (10 seconds), 68°C (20 seconds) and 72°C (20 seconds) for a total 10 cycles, before a final extension at 72°C (120 seconds). The product was purified using 0.6X AMPURE XP DNA purification beads, and the eluted PCR product was supplemented with nested FWR1-specific primers containing the read 1 site (H2F mix; 500nM), a tag-specific reverse primer (PT056; 500nM) and Q5 DNA polymerase to a final volume of 50µL, which then underwent PCR. The product was purified using (0.6X) AMPURE XP DNA purification beads, and a final PCR was performed to add Illumina indexes (500nM each), again using Q5 DNA polymerase to the amplified DNA. PCR product was gel extracted using the QIAquick gel extraction kit (QIAGEN) to ensure homogenous size selection, prior to size and concentration quantification using the Agilent TapeStation 4150 (Agilent Technologies) and the Qubit 2.0 Fluorimeter (Thermo Fisher). Libraries were submitted for sequencing using the MiSeq platform, via UCLGenomics, using the following read lengths 325bp, 8bp, 8bp and 275bp for read 1, the two index reads, and read 2, respectively.

### Bulk BCR raw sequence processing

Raw sequencing reads (fastq) were processed using the Immcantation framework via singularity (version 4.4.0) (Gupta et al., 2015; Vander Heiden et al., 2014). Briefly, reads were filtered (phred score > 20), prior to primer and UMI region identification used to define coarse read groups. These were then more finely clustered using cd-hit-est (Li and Godzik, 2006) with a threshold of 90%. Similarly to other published works, we assume that the resulting groups represent reads from unique RNA molecules (Galson et al., 2020). A consensus sequence was then generated per read group, and duplicate reads were collapsed to provide a single representative sequence. UMIs comprised of a single read were removed.

V, D and J regions were assigned using IgBLAST (Ye et al., 2013). Annotated sequences were then partitioned into coarse clusters based on identical V, J and CDRH3 lengths and then finely clustered based on 90% CDRH3 amino acid identity. Amino acid mutations were identified with shazam (Gupta et al., 2015).

For phylogenetic tree analysis, lineages containing multiple MBC subsets with at least 3 unique sequences were identified and processed using IgPhyML (Hoehn et al., 2019, 2017). CDRH3 sequence was ignored for this analysis as per the developer’s recommendations.

### Bulk BCR Bioinformatic analysis

Bulk BCR sequence analysis was carried out using R. Data manipulation and summarisation was primarily performed using the tidyverse suite of packages (Wickham et al., 2019), as well as foreach and doParallel for streamlining execution (Weston, 2022, 2022), and ape for basic analysis of trees (Emmanuel Paradis, Klaus Schliep, 2019). The vegan package was used to calculate Shannon and Simpson diversity indices (Jari Oksanen et al., 2024).

### De novo mutation calculation

Each lineage that contained multiple MBC subsets was first identified within each donor. Within these lineages, mutations were identified in BCR sequences by comparing each sequence to the lineage germline sequence. Next, BCR sequences underwent pairwise analysis to identify both shared and unique mutations for each sequence, and the proportion of unique mutations identified (unique mutations / total mutations). The comparisons were then grouped based on the subsets compared (e.g., TLM—Resting, TLM—TLM as multiple sequences from the same subset were also compared), and the median average taken, to provide a single value per comparison, per lineage. Once computed for all lineages, this was then summarised by taking the mean average per comparison, to give a single value per donor, which underwent statistical comparison.

### BCR phylogenetic network analysis

To ascertain the evolutionary ‘potential’ of the MBC subsets, phylogenetic networks were created, using igraph (Csardi and Nepusz, 2006). To gain evolutionary information, lineages containing multiple MBC subsets were first identified and phylogenetic trees estimated using IgPhyML. From these trees, distance between tips (sum of branch lengths) was calculated using the distTips function from the adephylo R package (Jombart et al., 2010), and assembled into a square matrix per lineage. To draw edges between related nodes, it was first necessary to define a distance threshold.

For this, the smallest tip-to-tip distance was identified for each tree, and a density curve created. For this curve, the lagged differences in y-axis value were divided by the lagged differences in x-axis, aiming to identify the position where the curve was most rapidly changing. This position was identified by taking the apex of the first identified peak (identified using the pracma package (Borchers, 2023)), and this x-axis position was the selected cut-off. By choosing the position where the aggregated tip-to-tip distances were most rapidly changing, we aimed to identify a relevant distance where BCRs were linked to neighbouring BCRs, and not linked to distant BCRs across multiple phylogenetic trees. This threshold was used to derive an adjacency matrix, whereby nodes that had a distance smaller than the calculated threshold were connected. The network was then directed by only allowing connections from nodes that had a smaller distance from the inferred germline node to those with a larger distance from the inferred germline node. Statistics were then calculated for each node in each network, namely the hub score (networkR; (Ekstrøm, 2019)) and the degree-ratio (igraph). Higher hub scores are achieved when a node has many outgoing connections and is further influenced by the quality of these downstream connections. Therefore, in our phylogenetic networks, nodes with higher hub scores reflect nodes with more close evolutionary relationships and indicates antigen driven SHM. The degree-ratio (number of outgoing connections divided by the number of incoming connections) was calculated using igraph and reflects the local environment of each node in the network. Statistical analysis was performed using Mann-Whitney U tests, via the ggpubr package (p > 0.05 not marked, p < 0.05 *, p < 0.01, **, p < 0.001 ***, p < 0.0001 ****) (Kassambara, 2023).

### Smart-Seq2 single-cell data analysis

Data were then processed using scanpy (v.1.9.8)(Wolf, Angerer et al. 2018) workflow with standard quality control steps; cells were filtered if number of genes >6000 or <600. Mitochondrial content was determined using scanpy.pp.calculate_qc_metrics function; cells with mitochondrial genes percentage <50% were retained for further analyses. Genes were retained if they were expressed by at least 2 cells. Gene counts for each cell were normalised to contain a total count equal to 10^6 counts per cell. This led to a working dataset of 301 cells. The top 2000 highly variable genes were selected based on Seurat v.3 algorithm (flavor = seurat_v3) with batch key “Sequencing_batch”. Highly variable genes were further refined by removing potentially confounding genes using the following search formula: ‘^HLA|^IG[HKL][VDJC]|^MT|^A[A-Z][0-9]|^B[A-Z][0-0]’. The number of principal components used for neighbourhood graph construction and dimensional reduction was set at 20. Data integration from both donors was performed using the bbknn algorithm^(Polanski, Young et al. 2020)^. Uniform Manifold Approximation and Projection (UMAP; v3.10.0)^(McInnes, Healy et al. 2018)^ was used for dimensional reduction and visualisation with all parameters as per default settings in scanpy. For the assessment of transcriptional similarity between Env-reactive B cells and MBC reference subsets, Celltypist^(Dominguez Conde, Xu et al. 2022)^ probability scores were generated as per default settings.

### 10X data analysis

#### Data alignment and quantification

Droplet libraries were processed using Cell Ranger v7.0.1; reads were aligned to the GRCh38 human genome (version refdata-gex-GRCh38-2020-A, 10X Genomics). CITE-seq UMIs were counted for GEX and ADT libraries simultaneously to generate feature-X droplet UMI count matrices.

#### CITE-seq data processing

Filtered and raw Cell Ranger output count matrices were integrated into a MuData object and further processed using muon package pipeline v0.1.5(Bredikhin, Kats et al. 2022). ADT counts for each protein were subjected to DBS (denoised and scaled by background) normalization with default settings. T/B cell doublets were removed using gene2filter if mutual expression at least two of the following markers was detected: CD19, CD4, CD8 (with positive expression thresholds set at 6, 12 and 12 for each respective marker).

#### Single-cell RNA seq quality control, normalization, embedding and clustering

Scrublet (v0.2.3)(Wolock, Lopez et al. 2019) was applied to each sample to generate a doublet score. This generated binomial distribution with an automated (default) threshold for doublet definition. Combined raw data from all samples were processed using scanpy (v1.9.8) workflow with standard quality control steps: cells were filtered if number of genes >5000 or <400. Cells with mitochondrial and ribosomal genes percentage <10% and >5%, respectively, were retained for further analyses (total 48,602 cells post QC) Genes were retained if they were expressed by at least 3 cells. Data were normalized to contain a total count equal to 10^4^ per cell and log + 1 corrected. Top 2000 highly variable genes (HVG) were selected based on Seurat v.3 algorithm (flavor = seurat_v3) with batch key “sample” and refined by removing the following genes ‘^HLA|^IG[HKL][VDJC]|^MT|^A[A-Z][0-9]|^B[A-Z][0-0]|XIST’. The number of principal components used for neighbourhood graph construction and dimensional reduction was set at 20, number of neighbours at 10. Uniform Manifold Approximation and Projection (UMAP; v3.10.0)^(McInnes, Healy et al. 2018)^ was used for dimensional reduction and visualisation with all parameters as per default settings in scanpy. Clustering was performed using the Leiden algorithm with an initial resolution of 0.4. CellTypist v1.6.2 (model: Immune_All_Low.pkl) with majority voting was used for assisted initial cell annotation. Top 10 differentially expressed genes were calculated using the Wilcoxon rank-sum test with minimum log fold change (LFC) set at 2. For detailed B cell analysis, non-B/plasma cells were filtered out and the remaining dataset re-processed from HVG selection step again with the same setting but with Leiden clustering resolution 0.6.

#### Differential abundance testing

The differential abundance of cell types across timepoints was determined using milopy (Milo framework) package (v0.1.0) standard workflow with default settings(Dann, Henderson et al. 2022).

#### Cell-cell interactions inference

Inferred number and strength of cell-cell interactions between all annotated cell subsets during pre-blip timepoint was calculated using CellChat v2(Jin, Plikus et al. 2024), following the standard workflow (from Anndata object) and keeping the default settings.

#### Gene set enrichment analysis and scanpy score

Gene set enrichment analysis (GSEA) was performed on selected MSigDB v7.5.1 Hallmark and KEGG genesets using the fgsea package (available on Bioconductor v3.18) in R (v4.3.3) and visualised with the GOChord function in the GOplot package. Briefly, genes were ranked in the descending order by the Wilcoxon statistic value from the pairwise Wilcoxon rank sum test (via ‘tl.rank_genes_groups’ in scanpy, comparisons: Blip vs. pre-Blip, Post-blip vs. Pre-blip). Expression scores for selected MSigDB v7.5.1 Hallmark & KEGG and GOBP genesets (GO:0050864, GO:0030888, GO:0045577) were generated by using using scanpy.tl.score_genes function.

#### Pseudotime analysis

GEX-based pseudotime and trajectory inference was performed on the whole B-cell dataset using slingshot package (available on Bioconductor v3.18)(Street, Risso et al. 2018) in R (v4.3.3) with default settings but specifying start.clust = ‘Naive1’. VDJ-based pseudotime inference was performed using ‘VDJ pseudobulk feature space’ workflow as outlined within dandelion package v0.3.3(Suo, Polanski et al. 2024).

#### BCR pre-processing

Single-cell V(D)J data from the 5′ Chromium 10x kit were initially processed with Cell Ranger multi pipeline (v7.0.1). BCR contigs contained in ‘filtered_contigs.fasta’ and ‘filtered_contig_annotations.csv’ from all samples were preprocessed using ‘immcantation’ inspired pre-processing pipeline(Suo, Polanski et al. 2024) implemented in the dandelion package v0.3.3 via Singularity following the standard workflow and recommended parametrization. This Singularity-based pre-processing also incorporates mutational burden quantification via SHazaM’s observedMutations(Gupta, Vander Heiden et al. 2015). Dandelion output files from each sample were then concatenated into a vdj object using dandelion function concat(). Contigs assigned to cells that passed QC on the transcriptome data were retained for further QC assessment using dandelion function pp.check_contigs()(Suo, Polanski et al. 2024).

#### B cell clone/clonotype analysis

BCRs were grouped into clones / clonotypes using dandelion package function dl.tl.find_clones(), which applies the following sequential criteria to both heavy- and light-chain contigs: (1) identical V and J gene usage, (2) identical junctional CDR3 amino acid length, and (3) at least 85% amino acid sequence similarity at the CDR3 junction (based on hamming distance). Light-chain pairing was performed using the same criteria within each heavy-chain clone**(Suo, Polanski et al. 2024)**. BCR networks were subsequently constructed using dandelion function: tl.generate_network(), which is based on adjacency matrices computed from pairwise Levenshtein distance of the full amino acid sequence alignment for BCR(s) contained in every pair of cells within each timepoint. Construction of the Levenshtein distance matrices were performed separately for heavy-chain and light-chain contigs, and the sum of the total edit distance across all layers/matrices was used as the final adjacency matrix as described previously**(Suo, Polanski et al. 2024)**. Weighted clonal overlap between each B cell subset at a single timepoint was calculated with dandelion function: tl.clone_overlap() with default settings**(Suo, Polanski et al. 2024)**. Clonality was assessed by calculating clone size Gini index after running dandelion function: tl_clone_diversity() with metric=”clone size”. This diversity metric does not rely on BCR network**(Suo, Polanski et al. 2024)**.

